# Parvalbumin neurons, temporal coding, and cortical noise in complex scene analysis

**DOI:** 10.1101/2021.09.11.459906

**Authors:** Jian Carlo Nocon, Howard J. Gritton, Nicholas M. James, Rebecca A. Mount, Zhili Qu, Xue Han, Kamal Sen

## Abstract

Cortical representations supporting many cognitive abilities emerge from underlying circuits comprised of several different cell types. However, cell type-specific contributions to rate and timing-based cortical coding are not well-understood. Here, we investigated the role of parvalbumin (PV) neurons in cortical complex scene analysis. Many complex scenes contain sensory stimuli which are highly dynamic in time and compete with stimuli at other spatial locations. PV neurons play a fundamental role in balancing excitation and inhibition in cortex and sculpting cortical temporal dynamics; yet their specific role in encoding complex scenes via timing-based coding, and the robustness of temporal representations to spatial competition, has not been investigated. Here, we address these questions in auditory cortex using a cocktail party-like paradigm, integrating electrophysiology, optogenetic manipulations, and a family of spike-distance metrics, to dissect PV neurons’ contributions towards rate and timing-based coding. We find that suppressing PV neurons degrades cortical discrimination of dynamic sounds in a cocktail party-like setting via changes in rapid temporal modulations in rate and spike timing, over a wide range of time-scales. Our findings suggest that PV neurons play a critical role in enhancing cortical temporal coding and reducing cortical noise, thereby improving representations of dynamic stimuli in complex scenes.

**Significance Statement:** One impressive example of sensory perception by the brain is its ability to analyze complex scenes, e.g., following what a friend is saying at a party amongst other speakers. Although some humans can solve this problem with relative ease, it remains very difficult for humans with a variety of impairments, e.g., hearing impairments, ADHD, and autism. The brain mechanisms underlying complex scene analysis remain poorly understood. Here, we recorded neural activity in auditory cortex in a complex auditory scene. When we suppressed PV neuron activity in auditory cortex, cortical performance decreased, and the timing of cortical responses was degraded. Our findings suggest that PV neurons improve the brain’s ability to analyze complex scenes by enhancing the timing of cortical responses while reducing cortical noise.

## Introduction

The cerebral cortex is critical for perception, attention, decision-making, memory, and motor output. Understanding the cortical circuit mechanisms that underly these functions remains a central problem in systems neuroscience. One line of investigation towards addressing this problem has been to identify the underpinnings of the cortical code; specifically, to assess whether cortical coding relies on rate or spike timing^1^. Previous studies have demonstrated both rate and spike timing-based coding in cortex^2–4^. However, a mechanistic understanding of how cortical circuits implement these codes and on what timescales is still missing. A second line of questioning towards addressing this central problem has been to utilize a combination of anatomy, physiology and optogenetics to interrogate cortical circuits and neuron types^5, 6^. This concerted approach has allowed systems neuroscience to identify key contributions of specific cell types to cortical circuits, including inhibitory neurons (e.g., parvalbumin-expressing (PV), somatostatin-expressing (SOM) and vasoactive intestinal peptide-expressing (VIP) neurons)^5^. However, the specific contributions of these diverse cell types to the cortical code remain unclear.

A potentially powerful strategy for unraveling cell type-specific contributions to cortical coding is to investigate problems where cortical processing is likely to play a central role. An important example of such a problem is complex scene analysis, e.g., recognizing objects in a scene cluttered with multiple objects at different spatial locations. The brain displays an astonishing ability to navigate such complex scenes in everyday settings, an impressive feat yet to be matched by state-of-the-art machines. The relative contribution of specific cell-types to this powerful computational ability remains unclear.

PV neurons are the most prominent group of inhibitory neurons in cortex^7^. Previous studies have investigated the role played by PV neurons in the generation of oscillations^8^ and spike synchronization^9^. PV neurons play a fundamental role in balancing excitation and inhibition^10^ and determining receptive field properties in cortex^11–13^. Optogenetic manipulation of PV neurons has provided insights into cortical responses, network dynamics, and behavior^14–19^. Specifically, a study by Moore et al.^19^ revealed that optogenetic suppression of PV neurons led to a rapid rebalancing of excitation and inhibition in cortex, with the expected increase in the activity of excitatory neurons, but a counterintuitive increase in the activity of inhibitory neurons. As elegantly dissected in the study, this occurs because the suppression of PV neurons leads to an increase in activity of excitatory neurons, which then drives both excitatory and inhibitory neurons downstream, rapidly rebalancing cortical activity. This result illuminates a property of cortical networks consistent with theoretical models but raises another question: does the suppression of PV neurons impact cortical temporal coding? The biophysical properties of PV neurons are well-suited for rapid temporal processing^5^ and therefore may be essential in the cortical temporal coding of dynamic stimuli present in complex scenes. Additionally, narrow-spiking units, which are thought to be putative inhibitory neurons, have exhibited distinct temporal response patterns to stimulus envelopes compared to those of regular-spiking units^20^. This motivates several open questions: do PV neurons play a critical role in cortical temporal coding of dynamic stimuli? Are such temporal codes robust to competing stimuli at other locations in space? Here, we address these questions in auditory cortex using a combination of electrophysiology, optogenetic suppression of PV neurons, and a family of spike distance metrics^21, 22^ to dissect specific contributions of PV neurons to the cortical code.

The auditory cortex (ACx) is well suited to investigate these issues. It is thought to play a key role in solving the cocktail party problem^23, 24^, one of the most impressive examples of complex scene analysis. Here, we integrate a cocktail party-like paradigm^25^ with optogenetic suppression of PV neurons to investigate the specific contribution of PV neurons to temporal coding in mouse ACx. We find that suppressing PV neurons degrades discrimination performance, specifically temporal coding, in ACx, and degrades performance over a wide range of time-scales. Our results reveal that despite the rebalancing of excitation and inhibition in cortical networks observed previously, suppression of PV neurons disrupts coding throughout ACx suggesting an important influence of PV neurons on cortical temporal coding and cortical noise.

## Results

We recorded single units (SUs) and multi-units (MUs) using a multielectrode array with 4 shanks and 32 channels throughout different layers in ACx of PV-Arch transgenic mice (Figures 1A-C and Supplementary Figure 1). We used a semi-automated detection and sorting algorithm to identify 124 units from *N* = 9 animals^26, 27^. Of these 124 units, 82 were identified as SUs (e.g., Figure 1C) while the remaining 42 were identified as MUs. In the results below, we focus on SUs. The results for MUs are given in Supplementary Figure 4C. Out of the SUs, 73 were identified as regular spiking (RS) while the remaining 9 were identified as narrow spiking (NS) based on the trough-peak interval of their mean waveforms (Supplementary Figure 2). RS and NS units have been found to correspond to excitatory and inhibitory neurons, respectively, in ACx^12, 28^.

**Figure 1.**
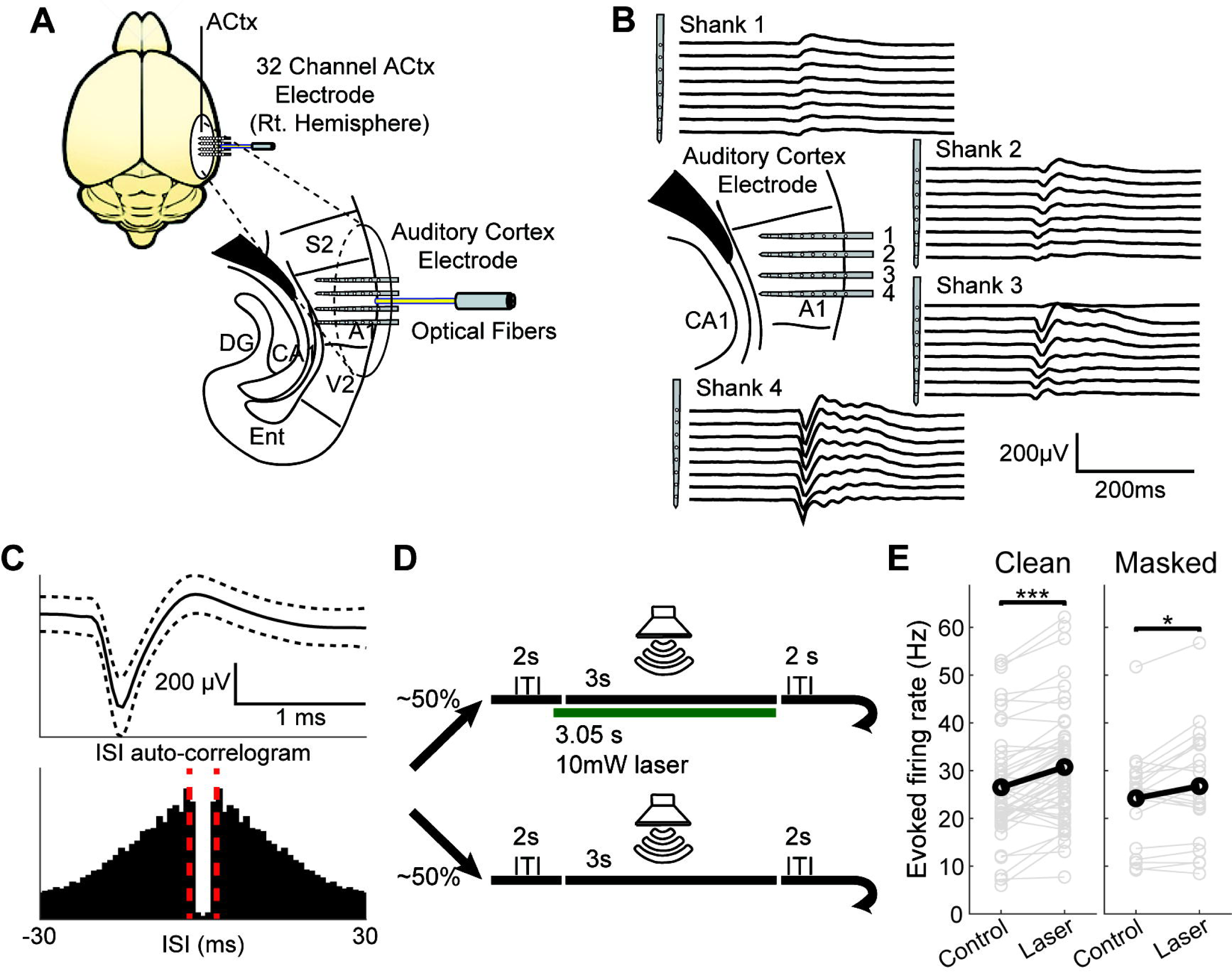
Experimental methods. **A:** Illustration depicting recording electrode location and optical fiber placement. Subjects were implanted with a 4-shank, 32-channel electrode array and optogenetic fiber in right hemisphere of ACx. Each shank contained 8 sites per shank with 100µm spacing between electrode contacts. **B:** Representative local field potential (LFP) activity from one mouse. LFP was used to estimate current source density and the layer of the recording site within each shank (Supplementary Figure 1). **C:** Example mean single unit waveform and inter-spike interval (ISI) auto-correlogram. Dashed lines in mean waveform represent one standard deviation above and below the mean, while scale bars are equal to 200µV and 1ms. Dashed red lines in correlogram represent ISIs of ±2ms. **D:** Schematic for control and optogenetic trial presentation. During approximately 50% of all trials, a 532nm laser would turn on 50ms before sound stimulus onset and turn off coincident with sound offset. **E:** Paired comparisons of mean evoked firing rate during control and laser trials. Paired *t*-tests yielded a significant increase in evoked firing rate during optogenetic suppression for clean (*n* = 49 configurations; *p* < 1e-04, *d* = −0.92) and masked trials (*n* = 20 configurations; *p* = 0.0219, *d* = −0.56).

To confirm specificity of expression, immunohistochemistry quantification was performed at the conclusion of the study and revealed that ∼93% of PV immuno-positive cells in auditory cortex were also Arch-GFP expressing neurons. Importantly, we also found that <1% of Arch-GFP expressing cells were immuno-negative for PV antibody (Supplementary Figure 3). Optogenetic suppression occurred on approximately 50% of trials, randomly interleaved, throughout the recording sessions. Suppression was achieved using light output from a 532nm laser that began 50ms prior to the auditory stimulus and consisted of continuous illuminations that co-terminated with sound offset. Within a given 800-trial session, optogenetic suppression strength remained constant (2mW, 5mW or 10mW), but was varied across sessions.

Next, we confirmed that the effects of optogenetic suppression of PV neurons in ACx were consistent with previous studies (Figure 1E, Supplementary Figures 4–5). Upon laser onset, we found that NS units in PV-Arch-expressing subjects showed an immediate suppression of spiking followed by an increase in activity (Supplemental Figure 4A), while NS units within non-Arch-expressing subjects did not show a change in activity during laser onset (Supplemental Figure 4B). We found that upon PV suppression, RS units increased their firing rate during both spontaneous and auditory evoked periods (Figure 1E and Supplementary Figure 5A), as expected^18, 19^. Different intensities enhanced the firing rate of RS neurons in a level-dependent manner consistent with previous studies (Supplementary Figure 5Ai). Counter-intuitively, but consistent with the previous study by Moore et al.^19^, NS units also increased their firing activity (Supplementary Figure 5B). As demonstrated by Moore et al. optogenetic suppression of PV neurons also produced a compensatory increase in inhibition and a rapid rebalancing of excitation and inhibition in cortex. Thus, the effects of optogenetic suppression of PV neurons on firing rates in ACx we observed are consistent with previous studies and the rapid rebalancing of excitation and inhibition. However, the effects of PV suppression on temporal coding in ACx remain unknown. Thus, we next inquired: does PV suppression impact temporal coding in ACx?

### Investigating cortical coding in mouse ACx using a cocktail party-like paradigm

To better understand cortical coding of complex scenes in a mouse model amenable to circuit interrogation using genetic tools, we adopted a cocktail party-like experimental paradigm^25^ while recording from neurons in ACx. Specifically, we recorded responses to spatially-distributed sound mixtures to determine how competing sound sources influence cortical coding of stimuli. The recording configuration consisted of 4 speakers arranged around the mouse at 4 locations on the azimuthal plane: directly in front (0°), two contralateral (45° and 90°) and 1 ipsilateral (−90°) to the right auditory cortex recording area. Target stimuli consisted of white noise modulated by human speech envelopes extracted from a speech corpus^29^. We utilized two target waveforms (target 1 and target 2) and a competing masker consisting of unmodulated white noise. Mice were exposed to either target-alone trials (Clean) or target-masker combinations (Masked) (Figures 2A-C).

**Figure 2.**
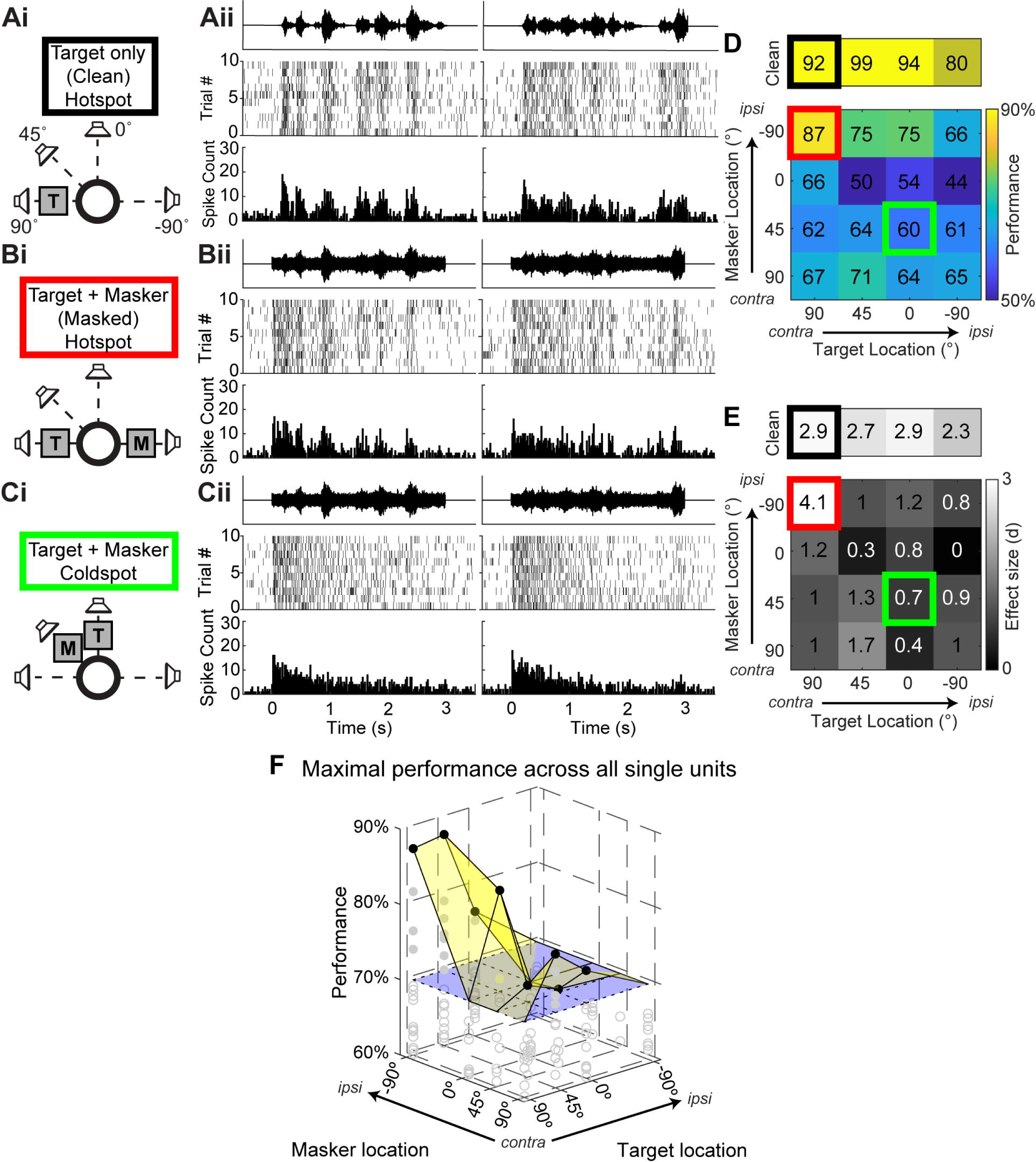
Cortical discrimination in a cocktail party paradigm in mouse ACx. **A:** Illustration depicting the stimulus configuration for clean trials originating at +90° azimuth (**Ai**) and responses to both target stimuli (T) (**Aii**). Auditory stimuli were presented from speakers at 4 locations. Target stimuli consisted of white noise modulated by human speech envelopes extracted from recordings of speech sentences (see Methods, Auditory stimuli). As shown in Aii, responses during clean trials exhibit spike timing and rapid firing rate modulation that follow the amplitude envelope of both target stimuli. All plotted PSTHs have a bin length of 20ms. **Bi:** Stimulus configuration for trials where targets (T) played at +90° and a competing masking stimulus (M) played at −90°. Masking stimuli consisted of unmodulated white noise with the same onset and offset times as target stimuli. **Bii**: Responses to masked trials shown in Di. In this configuration, spike timing and firing rate modulation follow both target stimuli, despite the presence of the competing masker. **Ci:** Stimulus configuration for trials where targets played at 0° and maskers played at +45°. **Cii**: Responses to target-masker configuration shown in Ci. For this configuration, spike timing and firing rate modulation do not follow either target stimulus, resulting in similar responses between target identities. **D:** Neural discriminability performance for all possible target-masker location configurations, referred to as the spatial grid, for the example cell featured in A-C. Outlined spots indicate configurations shown in A-C, matched by the outline color. Performance is calculated using a template-matching approach based on differences in instantaneous firing rate and spike timing similarities between spike trains (see Methods, Neural discriminability using SPIKE-distance). The top grid shows discriminability for clean trials on top, while the bottom grid shows discriminability for masked trials. All blocks are color-coded according to the color axis shown to the right of the masked grid. Configurations with high performance (2 70%) and a large effect size (d 2 1), e.g., the configurations outlined in black and red, are referred to as hotspots. **E:** Effect sizes for each spatial grid configuration in D, with the same outlines corresponding to examples in A-C. Positive values represent an increase in performance relative to a null distribution where spike trains within each target are template-matched to each other, while negative values represent a decrease in performance relative to null. **F:** The performance of all 23 single units exhibiting at least one hotspot during control trials. The translucent yellow surface represents the upper envelope of best performance across all single units for each masked spatial configuration, while the translucent blue surface represents the performance threshold of 70% for hotspots. Solid gray markers represent masked configurations with performances above threshold, while unfilled gray markers represent data points with performances below threshold. Black markers represent the maximal performance used to represent the upper envelope.

### Mouse ACx neurons show spatial configuration sensitivity between competing auditory stimuli

We assessed cortical coding using neural discriminability, which refers to the ability to determine stimulus identity based on neural responses and thus a neuron’s ability to encode stimulus features, and a variety of other quantitative response measures. Neural discriminability between the two targets (% correct) was computed both without the masker (Clean) (Figure 2A); and with the masker (Masked), for all possible combinations of target and masker locations (Figures 2B-C). We refer to the matrix of performance values from all speaker configurations as spatial grids, which allow for visualization of the spatial tuning sensitivity of a given unit in the presence of competing auditory stimuli. Values near 100% and 50% respectively represent perfect discriminability and chance discriminability, and positions of high performance (≥ 70%), which were also statistically significant (p ≤ 0.05) with a relatively large effect size (d ≥ 1), were deemed as hotspots. These hotspots represent locations of enhanced discriminability between the two targets, either in the absence (Clean) or presence (Masked) of a competing masking stimulus, using a spike distance-based classifier to determine how well target identity can be predicted given the spike train from that site based on dissimilarities in spike timing and instantaneous rate^25^ (Figure 2D: see Methods, Neural discriminability using SPIKE-distance).

Figure 2A illustrates spike trains from an example SU that shows high discriminability under both target-only conditions (Figures 2A, 2D, and 2E: black); and for a specific spatial configuration in the presence of a competing noise masker (Figures 2B, 2D, and 2E: red). In the masked condition, discriminability depends strongly on the spatial configuration of the target and masker, indicating that the response of this neuron is spatial configuration-sensitive (Figures 2C-E, red versus green).

Previous studies have demonstrated that neurons with the highest performance are most strongly correlated with behavior and strongly constrain population performance^30–34^. Thus, we were curious to test how the performance of the best neurons in our population would be affected by optogenetic suppression of PV neurons. To do so, we focused on the neurons with high discrimination performance in our population (i.e., SUs with at least one hotspot in the clean or masked conditions). 23 SUs showed hotspots at one or more spatial configurations, and there were 49 hotspots in the clean condition and 20 hotspots in the masked condition giving a total of 69 hotspots.

At each spatial configuration we observed a broad range of performance levels, consisting of neurons with significant hotspots (Figure 2F; filled circles), as well as neurons with poor performance (open circles), reflecting that different neurons in the population had different spatial configuration sensitivities. The upper envelope of maximal performance was relatively high for all spatial configurations, except co-located target-masker and ipsilateral target positions. Thus, as a population, ACx neurons showed robust performance at all spatial configurations in the contralateral hemisphere, when the target and masker were spatially separated. We did not observe any statistically significant differences in performance between SUs across different layers, or SUs with different waveform types (RS vs. NS) (Supplementary Figure 6).

### Suppression of PV neurons reduces discrimination performance at hotspots

To investigate the role of PV interneurons in auditory discrimination performance, we compared discrimination performance at hotspots, with and without optogenetic suppression of PV neurons in ACx. Figure 3A shows an example SU with and without suppression. Compared to the control response (Figure 3Ai), the optogenetic response (Figure 3Aii) shows an increase in spiking between the peaks of both target stimuli. Specifically, the responses exhibited an earlier onset and decreased spike timing reproducibility across trials during suppression (Figure 3Aiii). Figure 3B shows the spatial grids for the same example SU during both conditions, with the example configuration in Figure 3A (Clean Target 45°) outlined in black in the control grid (Figure 3Bi) and in red in the optogenetic grid (Figure 3Bii). This unit showed a decrease in performance across all clean configurations, and the hotspots in the control masked condition (Target 90°, Masker −90°; Target 45°, Masker −90°; Target 45°, Masker 90°) showed a reduction in performance to below threshold. Overall, we found that performance decreased significantly in both clean (p < 1e-04) and masked (p = 2e-04) conditions during suppression (Figure 3C), a decrease that was not significant in mice that did not express PV-Arch (Supplementary Figure 7).

**Figure 3.**
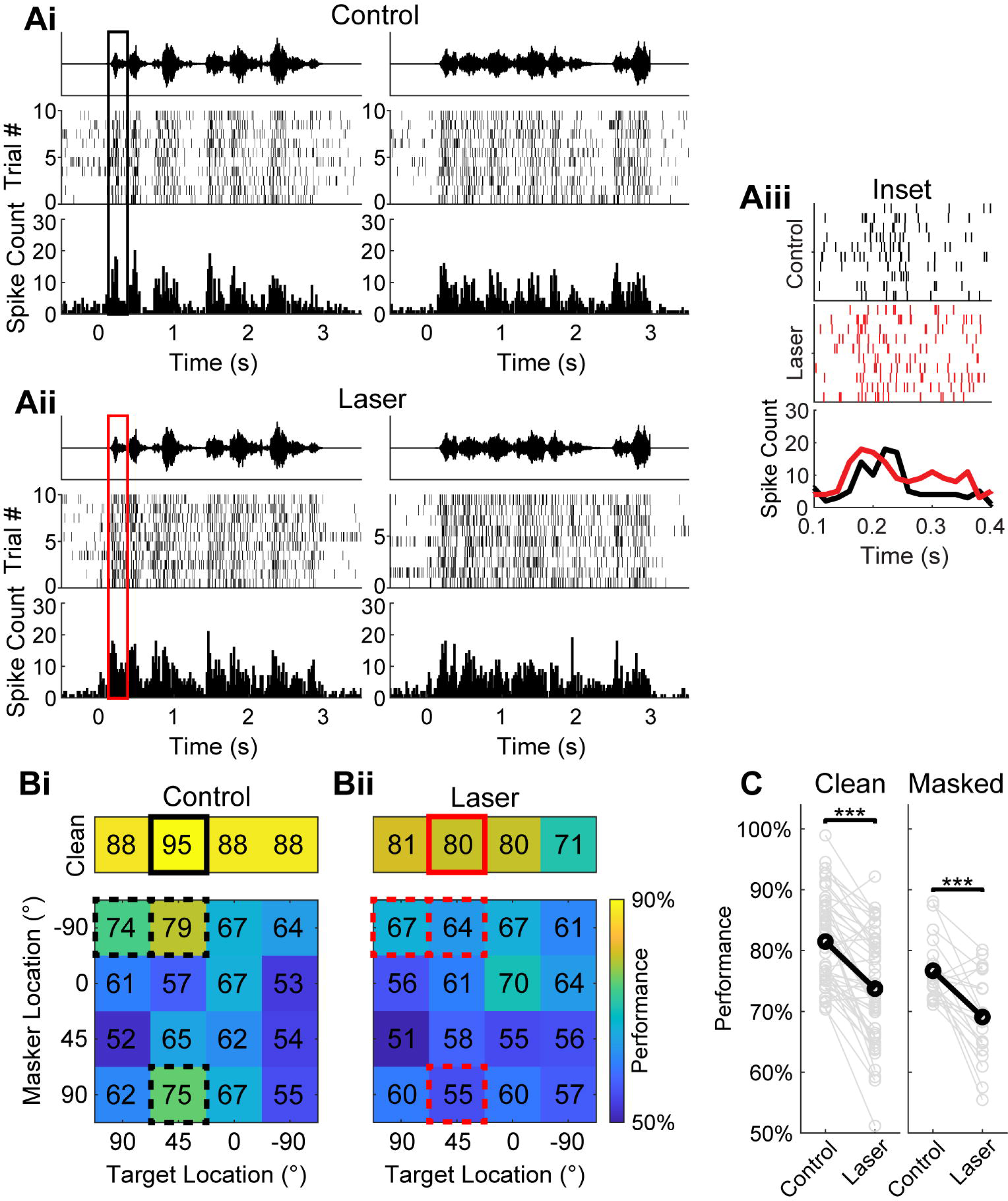
Effects of suppressing PV neurons on cortical discrimination. **A:** Responses during clean stimulus trials originating at 45° from one example cell during control (**Ai**) and optogenetic (**Aii**) conditions. **Aiii:** Inset showing zoomed-in portion of the response between 0.1 and 0.4s after sound onset, as outlined in Ai and Aii. Responses during optogenetic trials show earlier onset and reduced spike timing consistency, compared to the control. **B:** Example spatial grids from the same single unit during control (**Bi**) and optogenetic (**Bii**) conditions, with the performances at Clean Target 45° outlined in black to correspond to responses shown in A. Performance is color-coded according to the axis shown to the right of the Laser grid. The reduction in spike timing reproducibility during optogenetic suppression (seen in Aiii) contributes to the decrease in performance (80%) compared to control trials at the same configuration (95%). Additionally, performance decreased during optogenetic suppression for the rest of the clean configurations, while performance at the masked control hotspots, outlined by dashed boxes in both Bi and Bii, decreased to below threshold: Target 45°, Masker 90° (75% to 55%); Target 45°, Masker −90° (79% to 64%); and Target 90°, Masker −90° (74% to 67%). **C:** Paired comparisons of SPIKE-distance-based performance from control and PV-suppressed trials at the same spatial grid location. Paired *t*-tests yielded a significant decrease in performance for both clean (*n* = 49 configurations; *p* < 1e-04, *d* = 1.05) and masked (*n* = 20 configurations; *p* = 2e-04, *d* = 1.03) trials during optogenetic suppression, indicating that PV suppression significantly reduced discrimination performance.

### Suppression of PV neurons degrades cortical temporal coding

To determine the extent to which changes in the temporal dynamics of rapid firing rate modulation, spike timing, and average firing rate changes that occur during suppression might affect performance, we calculated different performance metrics across all hotspots. Specifically, we used inter-spike interval (ISI)-distance, rate-independent (RI)-SPIKE-distance, and spike count, as the basis for discriminability between spike trains. ISI-distance calculates the distance between two spike trains based on the dissimilarities in instantaneous firing rate modulation, while RI-SPIKE-distance measures spike timing dissimilarity between two trains while accounting for changes in firing rate differences^21^. Spike count distance is the absolute difference in the number of spikes between trains, effectively measuring differences in total firing rate. We found that performance based on both ISI-distance (Figure 4A) and RI-SPIKE-distance (Figure 4B) performances were relatively high. Both performances showed highly significant decreases with optogenetic suppression (ISI-distance-based performance: *p_clean_* < 1e-04, *p_masked_* = 0.0034; RI-SPIKE-distance-based performance: *p_clean_* < 1e-04, *p_masked_* = 0.0011). In contrast, performance based on spike count over the entire stimulus (Figure 4C) was close to chance level both for control and laser conditions, indicating that spike count alone was not sufficient to account for overall performance. The significant decrease in ISI distance-based performance indicates a disruption in rate-based coding, including the dynamics of instantaneous firing rate modulations. The significant decrease in RI-SPIKE based distance indicates that spike timing-based coding is also degraded by optogenetic suppression of PV neurons (Figure 4D).

**Figure 4.**
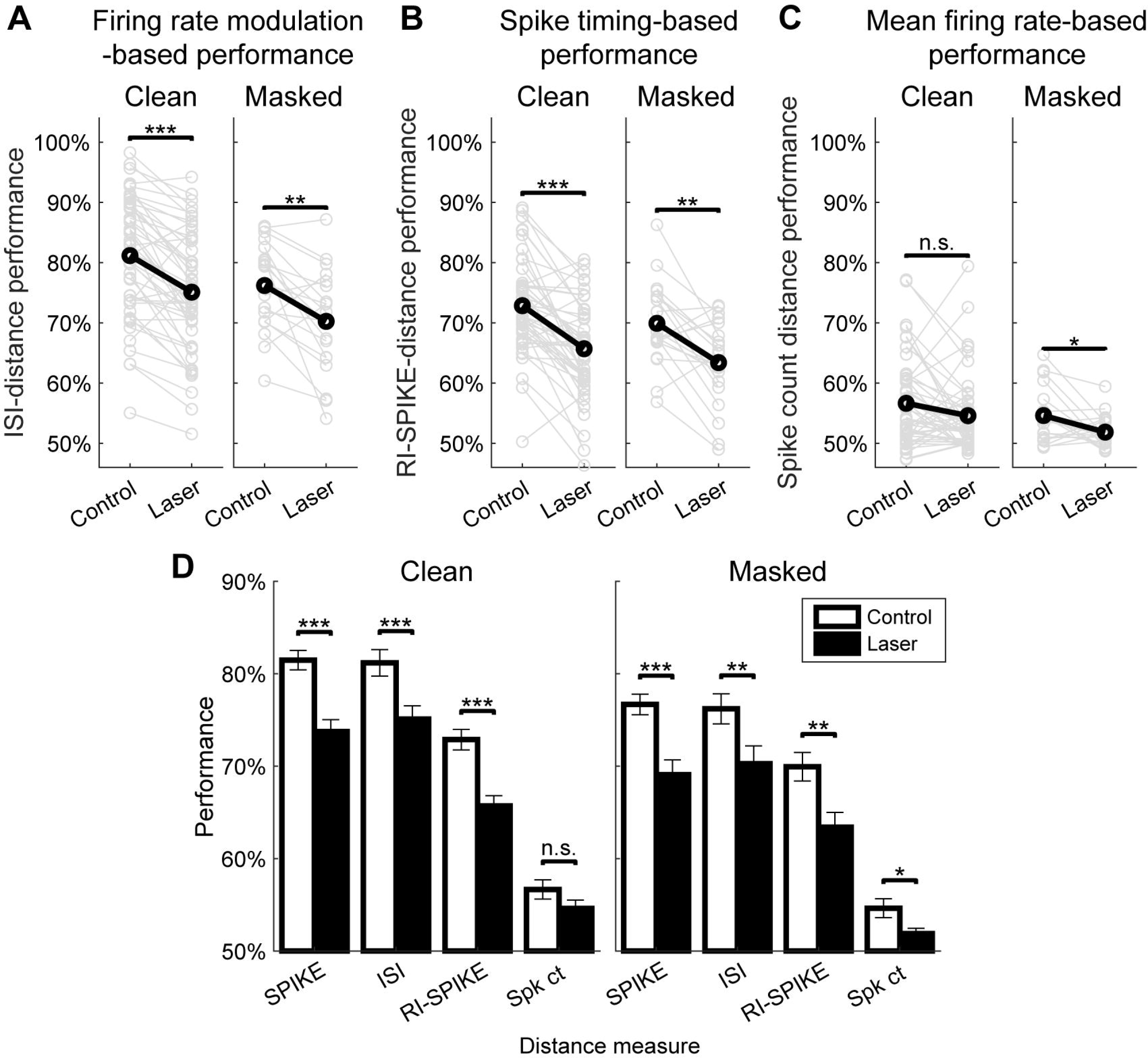
Effects of suppressing PV neurons on spike timing and rate-based coding measures. **A:** Performance based on ISI-distance, which measures differences between trains in instantaneous firing rate only (see Methods, ISI-distance). Paired *t*-tests showed a significant decrease in performance for both clean (*n* = 49 configurations; *p* < 1e-04, d = 0.92) and masked (*n* = 20 configurations; *p* = 0.0034, *d* = 0.75) trials. **B:** Performance based on RI-SPIKE-distance, which measures differences between trains in spike timing only (see Methods, RI-SPIKE-distance). Paired *t*-tests showed a significant decrease in performance for both clean (*p* < 1e-04, *d* = 0.95) and masked (*p* = 0.0011, *d* = 0.86) trials. **C:** Performance based on differences in total spike count between spike trains was near chance level, indicating that total spike count did not account for overall discrimination performance. Paired *t*-tests showed a significant decrease in performance for clean trials (*p* = 0.0590, *d* = 0.28) but not for masked trials (*p* = 0.020, *d* = 0.56). **D:** Summary figure showing contributions from spike distance measures presented in Figure 3C and 4A-C on the same scale and axis. Changes in spike timing and instantaneous firing rate-based measures (RI-SPIKE and ISI, respectively) provide relatively high discrimination performance and show a significant decrease upon optogenetic suppression of PV neurons.

### Effects on components of discrimination performance with suppression

Generally, discrimination performance depends on both the dissimilarity of responses between targets, as well as the similarity of responses within a target. To assess the relationship between different components of responses with performance, we calculated three metrics sensitive to firing rate and/or timing: the average firing rate; the rate-normalized root-mean-square (RMS) difference in the responses to the targets, which captures the difference in the temporal pattern of responses to the targets; and the trial similarity^35^, which captures the reproducibility of responses across trials within a target (see Methods).

We first calculated the correlation between evoked firing rate and performance during the control condition by pooling clean and masked data. Firing rate did not show a significant correlation with performance (*r* = −0.0645, *p* = 0.452), whereas both RMS difference and trial similarity measures were highly correlated with performance (rate-normalized RMS difference: *r* = 0.4205, *p* < 1e-04; trial similarity: *r* = 0.6013, *p* < 1e-04). These results suggest that both the pattern of firing rate modulations (quantified by RMS difference) as well as the reproducibility of responses (quantified by trial similarity) contribute to discrimination performance under control conditions. When comparing these measures between the control and laser conditions, we found that rate-normalized RMS difference significantly decreased with optogenetic suppression for both clean and masked trials (Figure 5A), and trial similarity significantly decreased during clean trials (Figure 5B).

**Figure 5.**
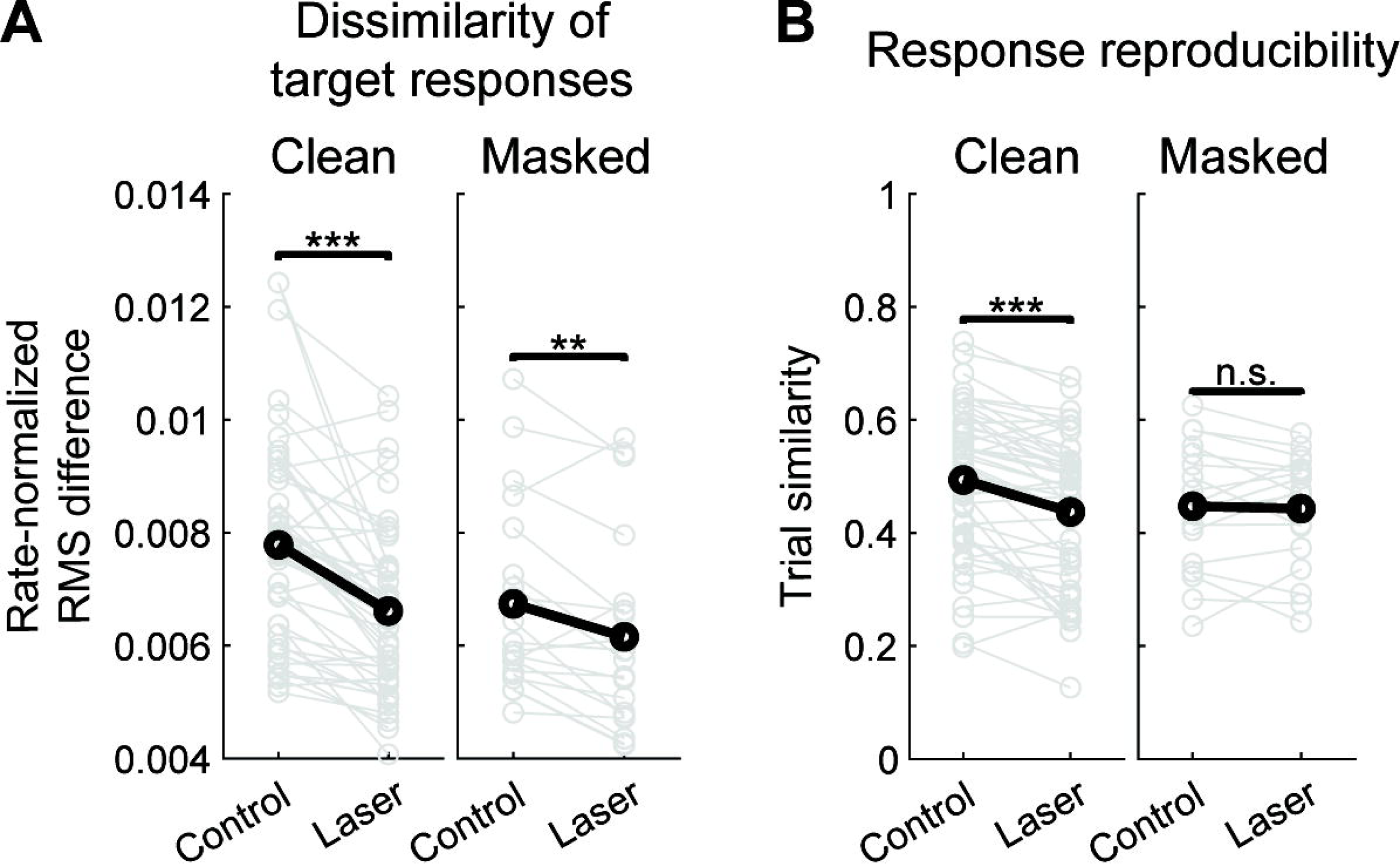
Effects of optogenetic suppression on spiking activity measures. **A:** Changes in dissimilarity of target responses via rate-normalized RMS difference between target PSTHs during both conditions. Paired *t*-tests found significant decreases between conditions during both clean trials (*n* = 49 configurations; *p* < 1e-04, *d* = 0.93) and masked trials (*n* = 20 configurations; *p* = 0.0031, *d* = 0.76). **B:** Changes in response reproducibility via trial similarity between responses to the same target during both conditions. Paired *t*-tests found a highly significant decrease between conditions during clean trials (*n* = 49 configurations; *p* < 1e-04, *d* = 0.85) but not masked trials (*n* = 20 configurations; *p* = 0.7333, *d* = 0.08).

### Optogenetic suppression decreases performance across a wide range of time scales

The previous analyses used spike distance measures which do not require a choice of a specific time-scale for analysis. A further interesting question regarding discrimination performance is the optimal time-scale for discrimination. Thus, we next quantified the time-scale for optimal discrimination using the van Rossum spike distance measure^36^ (see Methods). We found that optimal time-scales for discrimination (*τ*) for most neurons was around 40 milliseconds, with a significant proportion of neurons covering even finer time scales down to ∼10 ms (Figure 6A). Optimal *τ* was not significantly different between control and laser conditions for clean trials (*p* = 0.4920) but significant for masked trials (*p* = 0.0098), and performance decreased significantly in the laser condition across a wide range of time-scales (Figures 6B-D, Table 1). These results indicate that PV suppression did not significantly change the optimal time-scale for discrimination but rather degraded discrimination across a wide range of time-scales.

**Figure 6.**
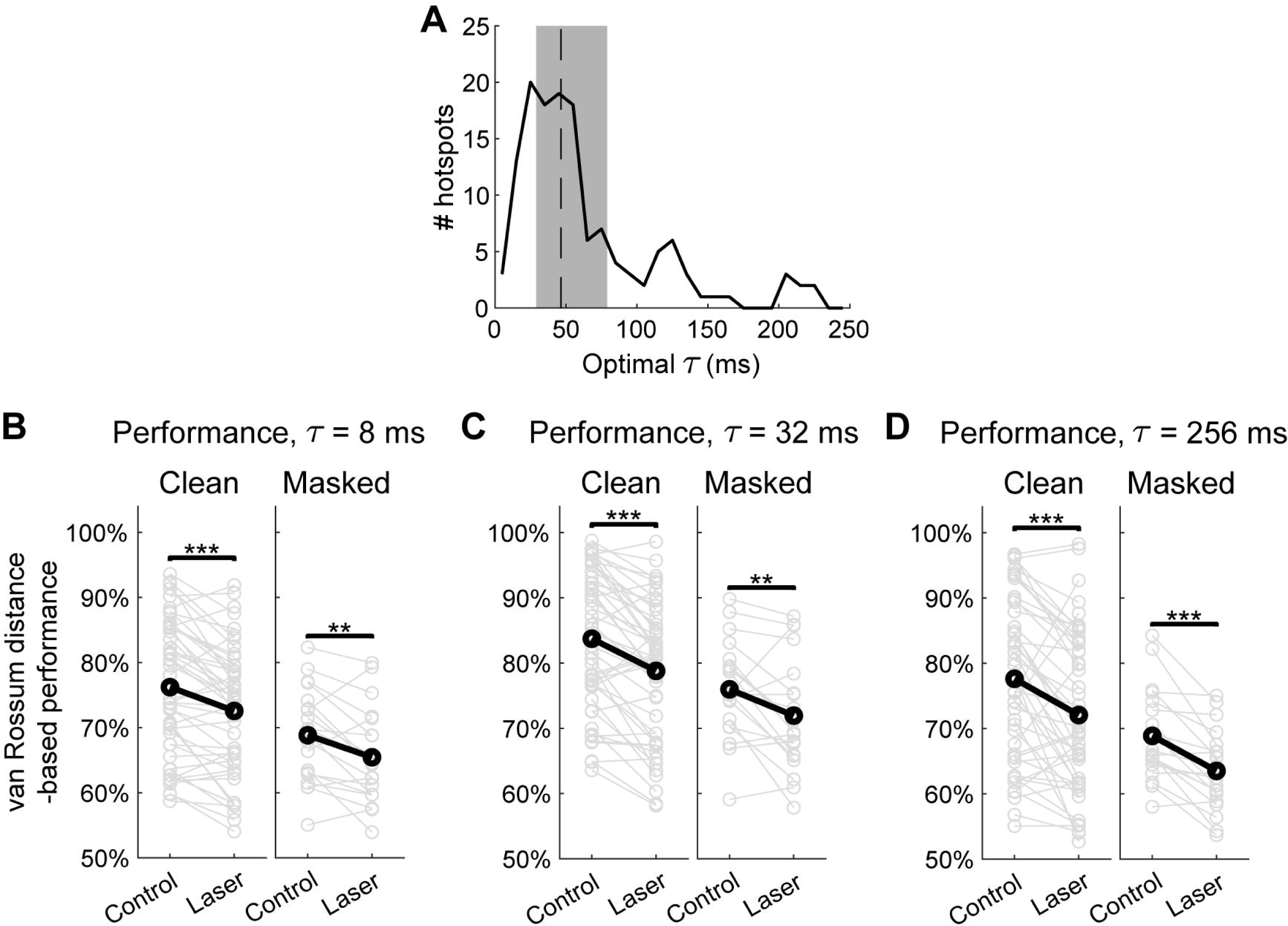
Decoding time analysis. **A:** Histogram of optimal τ for hotspots across both conditions (control and laser) and stimulus types (clean and masked). Dashed line indicates median value of 46.5ms, and shaded region represents the inter-quartile range (IQR) between 29ms and 79ms. Paired *t*-tests did not find a significant change in optimal *τ* within hotspots between conditions during clean trials (*n* = 49 configurations; *p* = 0.492, *d* = −0.10) but found a significant decrease during masked trials (*n* = 20 configurations; *p* = 0.0098, *d* = 0.64). **B:** van Rossum-based performance with τ set at 8ms. Performance was found to significantly decrease during both clean (*p* < 1e-04, *d* = 0.74) and masked (*p* = 0.0042, *d* = 0.73) trials. **C:** van Rossum-based performance with 1 set at 32ms. Performance was found to significantly decrease during both clean (*p* < 1e-04, *d* = 0.87) and masked (*p* = 0.0098, *d* = 0.76) trials. **D:** van Rossum-based performance with *1* set at 256ms. Performance was found to significantly decrease during both clean (*p* < 1e-04, *d* = 0.64) and masked (*p* < 1e-04, *d* = 1.18) trials.

**Table 1.** Effect sizes and paired *t*-test results for all *1* values used in van Rossum distance-based performance calculations.

| $\tau$ (ms) | $d_{clean}$ | $p_{clean}$ | $d_{masked}$ | $p_{masked}$ |
| --- | --- | --- | --- | --- |
| 1 | -0.02 | 0.9157 | 0.56 | 0.0226 |
| 2 | 0.11 | 0.4530 | 0.67 | 0.0074 |
| 4 | 0.35 | 0.0189 | 0.70 | 0.0055 |
| 8 | 0.74 | < 1e-04 | 0.73 | 0.0042 |
| 16 | 0.87 | < 1e-04 | 0.66 | 0.0079 |
| 32 | 0.87 | < 1e-04 | 0.64 | 0.0098 |
| 64 | 0.79 | < 1e-04 | 0.74 | 0.0035 |
| 128 | 0.72 | < 1e-04 | 1.03 | 2e-04 |
| 256 | 0.64 | < 1e-04 | 1.18 | < 1e-04 |

## Discussion

### Diversity of cortical cell types and the cortical code

One of the most striking features of the cerebral cortex is the tremendous diversity of its cell types^37^. Understanding the computational role of such diversity in cortical coding is central to systems neuroscience. Addressing this central question requires understanding cell type-specific contributions to the cortical code at both the single neuron and the population level. A small number of previous studies have demonstrated a role of specific cell types in cortical population coding; specifically, the generation of oscillations^38, 39^, and synchrony across cortical layers and areas^40, 41^. However, cell type-specific contributions to the cortical code at the single unit level, a fundamental aspect of cortical encoding, remain poorly understood. In this study, we addressed this fundamental gap by investigating the role of PV neurons in cortical coding of a complex scene, i.e., a cocktail party-like setting, in mouse ACx.

### Cortical discrimination & PV neurons; rate, spike timing and temporal codes

We assessed cortical coding using neural discrimination performance and other quantitative measures. There is a rich history of quantitative work on cortical discrimination^30, 32^. These studies have suggested a critical role for neurons with the highest levels of performance in a population, which correlate strongly with behavioral performance and determine the overall performance at the population level. In this study we examined cortical discrimination of dynamic stimuli in a complex scene by the highest performing neurons in ACx, extending the previous body of work in several ways: First, we assessed the impact of optogenetic suppression of PV neurons on discrimination performance. Optogenetically suppressing PV neurons resulted in increased firing rate during spontaneous and auditory evoked activity, which is consistent with the effects of inhibitory blockade on cortical responses^42–44^. A recent study by Moore et al.^19^ employed optogenetic suppression of PV neurons to powerfully reveal an important property of cortical networks: rapid rebalancing of excitation and inhibition upon PV suppression. Our study reveals that despite such rebalancing, cortical discrimination performance is degraded across cortical layers in ACx upon PV suppression. This finding suggests that PV neurons play a role in improving discrimination of dynamic stimuli in ACx, both sounds in quiet backgrounds, as well as in the presence of competing sounds from other spatial locations. Second, we quantified the contributions of instantaneous firing rate modulations, spike timing, and spike count towards cortical discrimination, using a family of spike distance metrics. These metrics provide a powerful set of tools for dissecting different components of cortical coding. Although these metrics have been employed in previous theoretical studies, to our knowledge this is the first time they were applied to analyze cortical responses. This analysis revealed that high discrimination performance is mediated by the temporal pattern of firing rate modulations and spike timing reproducibility, and that optogenetic suppression of PV neurons degraded both components.

Previous studies have demonstrated that auditory cortical neurons can employ both rate and spike timing-based codes^3, 4^; and provided insight into the roles of inhibitory neurons in shaping frequency tuning^11–13, 45^, frequency discrimination^14, 46^, adaptation^15^, sparseness^47^, and gap encoding^28^. An influential review on neural coding also defined a precise notion of a temporal code as one that contains information in spike timing beyond rate modulations^48^. Temporal codes have been challenging to identify because the contributions of rate vs. spike timing are often difficult to decouple. Our results based on spike distance measures, which quantify both rate-dependent and rate-independent components of coding, suggest that PV neurons specifically contribute to temporal coding in cortical discrimination. This computational portrait of PV neurons validates the importance of their established electrophysiological specializations–namely, fast, efficient, and temporally precise responses^5^.

### A general cortical representation for the cocktail party problem across species

From a comparative standpoint, we found that the key features present at the cortical level within ACx of the mammalian mouse was consistent with previous findings in songbirds. Specifically, a previous study by Maddox et al. found hotspots at particular spatial configurations of target and masker on the spatial grids of cortical level neurons in songbirds^25^. Songbirds and mice have different frequency ranges of hearing and therefore the cues used for spatial processing, e.g., interaural time difference (ITD), and interaural level difference (ILD) are frequency-dependent, and the peripheral representations of these cues are likely to be different across species with different frequency ranges of hearing. This suggests the emergence of general cortical representations for solving the cocktail party problem despite different peripheral representations of acoustic cues across species.

### Time scales of cortical discrimination

A further interesting question regarding cortical discrimination is: what is the optimal time scale for maximal discrimination performance? One characteristic time scale in our stimuli arises from the slow modulation of speech envelopes on relatively long time-scales ∼100-500 ms, or equivalently, in the 2-10 Hz frequency range^49^. We found that the optimal time-scale for most neurons in our dataset was much finer ∼40 ms, with a significant number at even finer timescales down to ∼10 ms. These time-scales are well matched to the duration of short ultrasonic vocalizations in mice (Figure 7), and finer grain structures within these vocalizations, e.g., spectral features and frequency sweeps^50^. These time-scales are similar to those found in a previous study of decoding sinusoidally amplitude-modulated (SAM) tones in mouse auditory cortex^35^, and consistent with integration time scales in cat auditory cortex^51^. Phonemic structures in speech also occupy similar time-scales, which are in the beta and low gamma range of frequencies^52, 53^. Thus, the time-scales for optimal discrimination in ACx, may be well suited for analyzing such vocalizations and the finer spectro-temporal features within.

**Figure 7.**
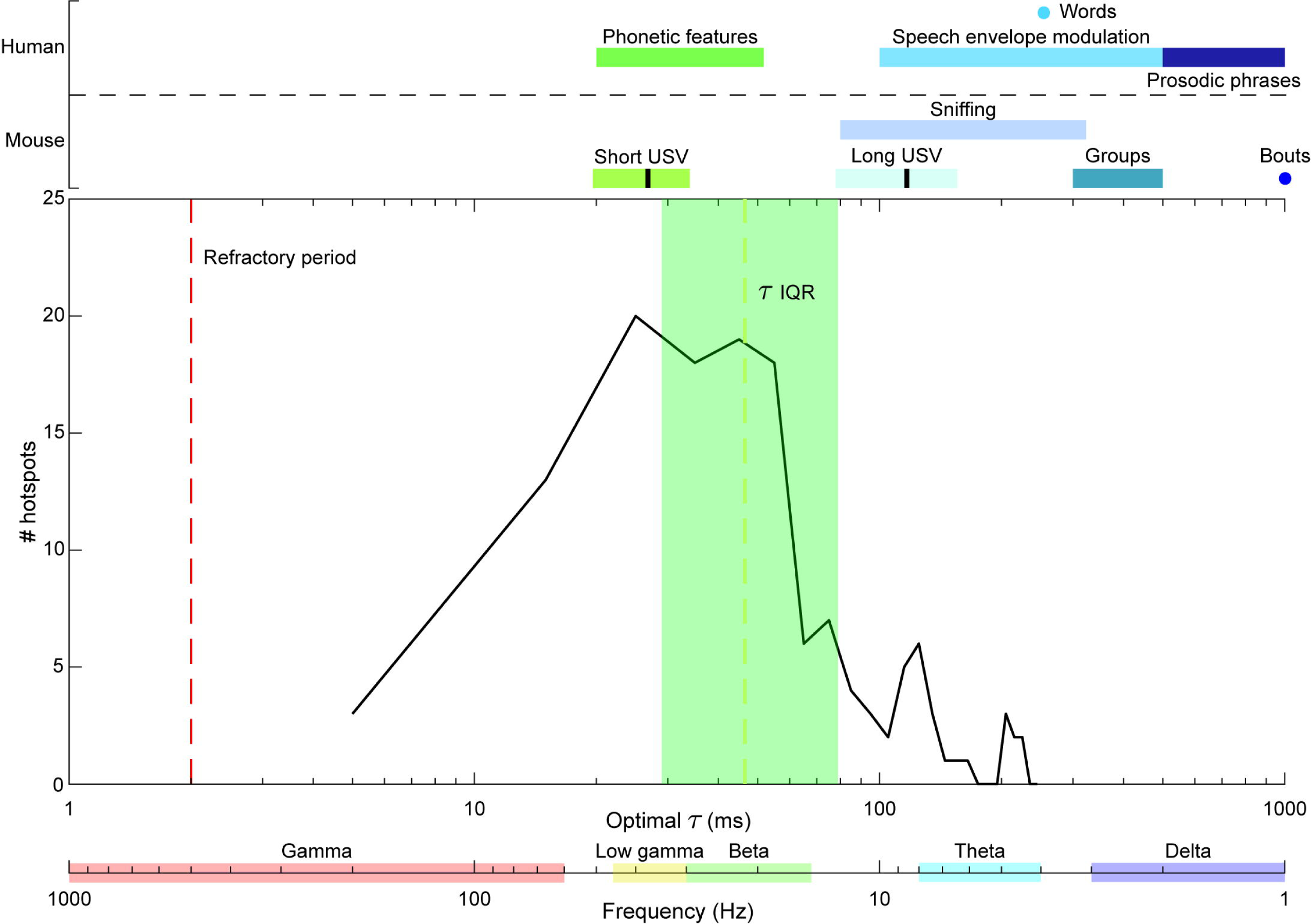
Comparison of optimal 1 values and other time scales. Semi-logarithmic plot showing various time-scales for spike timing in mouse ACx neurons compared to mouse vocalizations, human speech, and neural oscillations. Top: Time scales for human speech sounds, mouse vocalizations, and sniffing periods^74^. In the mouse time-scales, short and long USV bars represent the mean (black line) ± 2 SD. vocalization length. Within the plot, from left to right: the refractory period for single units (red dashed line); and the distribution of optimal *1* values from Figure 6 (solid black curve), with the dashed yellow line indicating the median value of 46.5ms and the shaded green region representing the IQR. The bottom axes show the time scale of optimal *1* values in ms and frequency in Hz, with the latter decreasing from left to right. Shaded bars represent frequency bands for neural oscillations.

### Cortical noise

Our findings are also relevant in the context of cortical noise, which can have a profound impact on cortical codes^54^. We found that suppressing PV neurons did not change the optimal time-scale for discrimination, but rather degraded performance at a wide range of time-scales (Figure 6). Additionally, we observed that suppression impacted specific components underlying discrimination: Most notably, PV suppression decreased the difference in the pattern of responses (quantified by RMS difference) between targets as well as decreasing the reproducibility of responses across trials (quantified by trial similarity). Taken together, these observations are consistent with an overall enhancement in cortical noise level across multiple time-scales upon PV suppression. A previous study on PV suppression in ACx observed the rapid rebalancing of excitation and inhibition, suggesting maintenance of the stability of global cortical representations^19^. However, our results suggest that despite this excitatory-inhibitory rebalancing, PV suppression also leads to an increase in cortical noise, fundamentally impacting the fidelity of cortical coding, including temporal coding.

### Within channel vs. cross-channel inhibition

Cortical inhibitory neurons can mediate feedforward, recurrent and di-synaptic feedback inhibition in cortical circuits, illustrated in a schematic model in Figure 8. Previous modeling studies have demonstrated that feed-forward within-channel inhibition can improve discrimination performance^55^; whereas inhibition across different channels can lead to the formation of hotspots and the specific pattern of spatial configuration sensitivity^56^. Our results suggest that PV neurons mediate within-channel inhibition, corresponding to I neurons in the schematic model. This is consistent with our finding that although suppressing PV neurons reduced discrimination performance, it did not completely eliminate the presence of hotspots on the spatial grids, suggesting that PV neurons alone do not control the emergence of hotspots. Based on these observations, we hypothesize that a separate cell-type (X neurons in Figure 8) mediates cross-channel inhibition, resulting in the generation of hotspots and the specific pattern of spatial configuration sensitivity on spatial grids. A candidate cell type that may correspond to X cells are somatostatin-positive (SOM) neurons, which have been implicated in di-synaptic feedback inhibition^57, 58^ and surround suppression^59, 60^. These distinct roles may be functionally well suited for solving the cocktail party problem, with one class of neurons (PV) enhancing the temporal coding of dynamic stimuli at a target location, and another class of inhibitory neurons (X) suppressing competing stimuli from other spatial locations.

**Figure 8.**
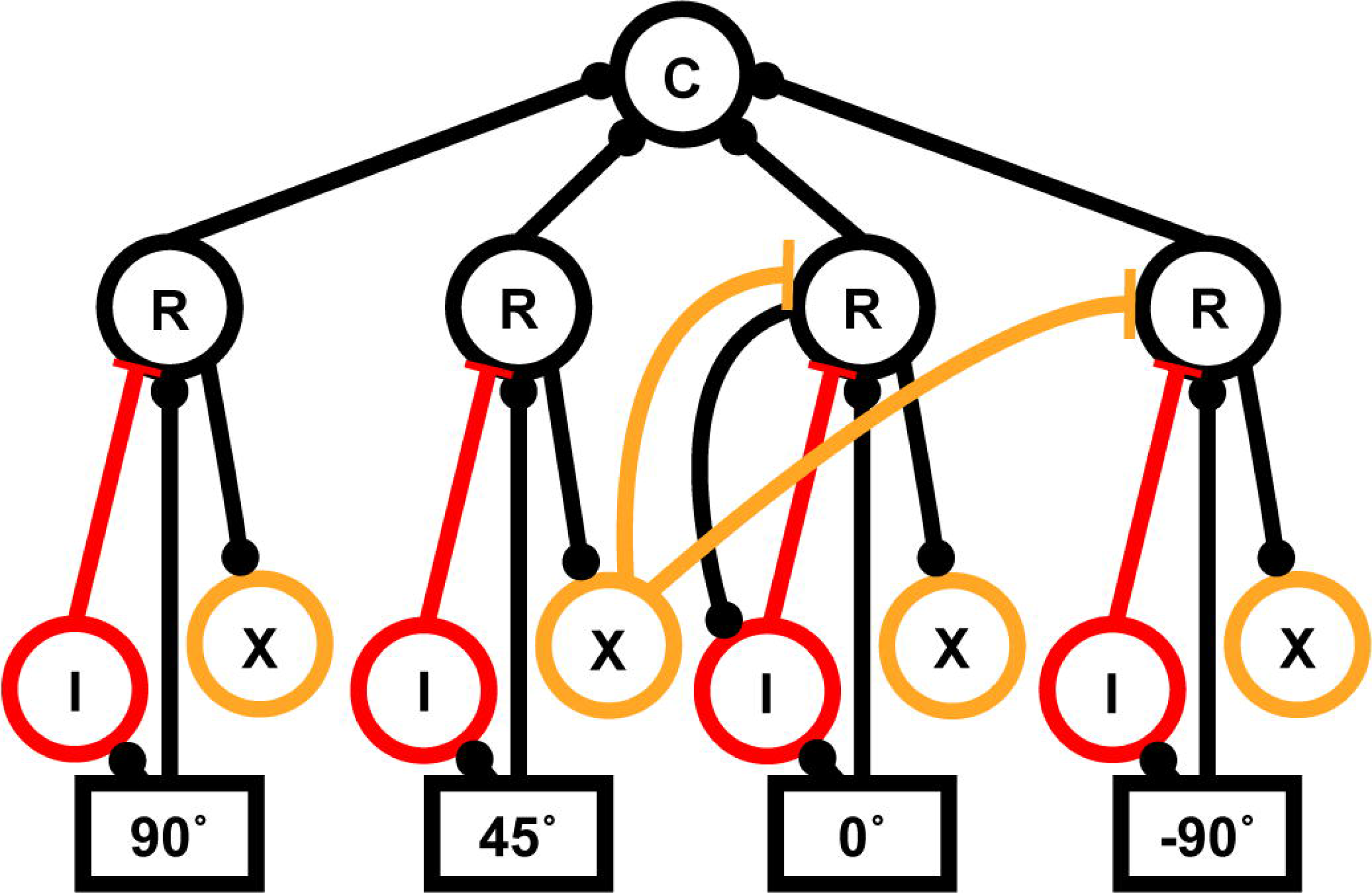
Cortical circuits for complex scene analysis. Hypothesized conceptual model of cortical circuit underlying spatial grids. C and R cells represent excitatory units, I cells mediate within-channel inhibition, and X cells mediate cross-channel inhibition.

### Limitations and future directions

Several limitations in this study should be further addressed in the future. Although we used a cocktail party-like paradigm to probe auditory cortical responses to dynamic stimuli our experimental paradigm had some limitations. First, the target stimuli did not have any specific behavioral significance, unlike the case of speech recognition at a cocktail party. Second, the masker stimuli did not contain any temporal modulations, unlike competing speakers at a cocktail party. Despite the anthropomorphic nature of our stimuli, we have demonstrated for the first time that auditory cortical neurons in mice are able to encode the distinct temporal features of both targets in the presence of a competing noise masker from different spatial locations. Future studies should address these limitations, e.g., by employing mouse communication sounds as targets and maskers. Although we were able to characterize the time-scale for optimal discrimination in ACx, we did not characterize the integration window, or the encoding window^48, 51, 61^. Future studies that characterize both the time-scale for optimal discrimination as well as the encoding window can address whether cortical neurons also employ temporal encoding, i.e., encode information in the temporal pattern of spikes within the encoding window^48^. Within this study, mice were awake but listening passively, whereas listening in a cocktail party-type setting is an active sensing process. It will be interesting to probe cortical coding in awake, behaving mice in experiments where animals attend to a specific spatial location. A recent theoretical model of attention in auditory cortex, the AIM network model^62^, suggests distinct roles for different interneuron groups in attentional sharpening of both spatial and frequency tuning which enables flexible listening in cocktail party-like settings, e.g., monitoring the entire scene, selecting a speaker at a spatial location, and switching to a speaker at a different location. Future experiments probing distinct interneuron populations (e.g., PV, SOM and VIP neurons) in behaving animals, in conjunction with testing and extending the AIM model may further unravel cortical circuits for solving the cocktail party problem.

## Materials and Methods

### Subjects

All procedures involving animals were approved by the Boston University Institutional Animal Care and Use Committee and the University of Illinois at Urbana-Champaign Institutional Animal Care and Use Committee (IACUC). A total of 14 mice were used in this study. Original breeding pairs of parvalbumin-Cre (PV-Cre: B6;129P2-Pvalbtm1(cre)Arbr/J), and Ai40 mice (Arch: B6.Cg-*Gt(ROSA)26Sor^tm40.1(CAG–aop3/EGFP)Hze^*/J) mice were obtained from Jackson Laboratory (Maine), and all breeding was done in house. Subjects consisted of both male and female PV-Arch (*n* = 9) offspring and PV-Cre (*n* = 5) only offspring (controls) 8-12 weeks old on the day of recording.

### Surgery

Mice were surgically implanted with a head-plate as described previously^63, 64^. Briefly, under isoflurane anesthesia, stereotaxic surgery was performed to install a headplate, electrode, and optical fiber. The custom head-plate was mounted anterior to bregma allowing access to ACx caudally. The headplate was anchored to the skull with 3 stainless steel screws and dental cement. A fourth screw was connected to a metal pin and placed in the skull above the contralateral cerebellum to serve as the reference. A craniotomy was made above the right auditory cortex (AP −2.3 to −3.6, ML + 4.0 to +4.5, DV). Using a stereotaxic arm, a 32-contact linear probe (Neuronexus, Ann Arbor, MI; model: a 4×8-5mm-100-400-177-CM32) with 100μm spacing between electrode contacts and 400μm spacing between shanks, was positioned into ACx, perpendicular to the cortical surface. Because of the curvature of the ACx surface, not all four shanks could be placed at precisely the same depth during each experiment. Probes were advanced until all electrode contacts were within the cortical tissue and shanks were positioned along the rostro-caudal axis of ACx (Figures 1A-C). An optical fiber, 200μm in diameter, was placed medially to the 4 shanks and positioned between the two innermost shanks terminating at the cortical surface (Figure 1A). After implantation, mice were allowed to recover for 4-7 days before undergoing habituation to being head-fixed as described below.

### Habituation

Following surgery and complete recovery, mice were first handled for several days before being head-fixed to the recording apparatus. Mice were gradually exposed to longer restraint periods at the same time of day as subsequent recording sessions. Each animal received at least 6 habituation sessions prior to the first recording day. Under head-fixed conditions, mice were loosely covered with a piece of lab tissue taped down on either side (Kimwipes: Kimberly-Clark, Irving, TX) to encourage reduced movement. At the conclusion of habituation, mice underwent recording sessions in the presence in the spatial stimuli as described below.

### Recording sessions and data acquisition

All recordings were made with a Tucker Davis Technologies (TDT; Alachua, FL) RZ2 recording system in an electrically-shielded sound attenuation chamber. Broadband neural signals at 24414.0625 Hz were recorded for each of the 32 channels. Local field potentials (LFPs) were band-pass filtered between 1 and 300 Hz, notch-filtered at 60 Hz, and digitized at 3051.8 Hz and used for current source density analysis (see Supplemental Methods).

Recording sessions consisted of both non-optogenetic and optogenetic trials in random order. The inter-trial interval was 5 seconds, with 3s of stimulus playback followed by 2s of silence. Mice were exposed to target-alone (clean) trials and target-masker (masked) combinations. 10 trials were given per target identity for all possible combinations of target location, masker location (including clean trials), and optogenetic suppression of PV neurons. Thus, animals received a total of 800 trials per ∼60 minute recording session, with each session having a set laser power.

### Auditory stimuli

All auditory stimuli were generated in Matlab and consisted of either target, masker, or combination of the two stimuli played from four TDT ES-1 electrostatic speakers. Target stimuli consisted of white noise modulated in time by human speech envelopes taken from the Harvard IEEE speech corpus^29^ which has been used in previous psychological studies of the cocktail party effect^65^. Masker stimuli consisted of 10 unique tokens of unmodulated white noise. Before speaker calibration, all stimuli were generated with the same RMS value, and sampling frequency was 195312 Hz to capture the frequency range of hearing in mice. Stimuli were loaded onto a custom RPvdsEx circuit on an RZ6 Multi I/O processor, which was connected to two PM2R multiplexers that controlled the location of target and masker stimuli during trials.

During recordings, the stimuli were presented 18 cm from the mouse’s head using four speakers driven by two TDT ED-1 speaker drivers. The four speakers were arranged around the mouse at four locations on the azimuthal plane: directly in front (0°), two contralateral (45° and 90°) and 1 ipsilateral (−90°) to the right auditory cortex recording area. Before recording sessions, stimulus intensity was calibrated using a conditioning amplifier and microphone (Brüel and Kjær, Nærum, Denmark; amplifier model: 2690, and microphone model: 4939-A-011). For 7 of the 9 Arch mice and the 5 PV-only control animals, all stimuli were at a measured 75dB intensity at the mouse’s head. For the remaining 2 Arch mice, stimulus intensity was set to 70dB. Stimulus playback lasted 3s with a 1ms cosine ramp at onset and offset.

### Optogenetic stimulation

Laser light for optogenetic stimulation of auditory cortex was delivered through a multimode optically-shielded 200µm fiber (Thorlabs, Newton, NJ; model: BFH48-200), coupled to a 532nm DPSS laser (Shanghai Laser Ltd., Shanghai, China; model: BL532T3-200FC), with the fiber tip positioned right above the cortical surface. Laser power was calibrated to 2mW, 5mW, or 10mW at the fiber tip using a light meter calibrated for 532nm wavelength (PM100D, Thorlabs, Newton, NJ). The intensity was determined based on optogenetic cortical PV suppression studies using Archaerhodopsin from the literature^14, 66^. During optogenetic trials, the laser was turned on 50ms before stimulus onset and co-terminated with the end of the auditory stimuli (Figure 1D). Square light pulses lasting 3.05s were delivered via TTL trigger from the RZ2 recording system to the laser diode controller (ADR-1805). Optogenetic trials were randomized throughout the recording session such that animals received all stimulus/masker pairs from each location with and without laser. Recordings were done in successive blocks with constant optogenetic suppression strengths of 2mW, 5mW, or 10mW, with each block lasting ∼60 minutes and having their own set of control trials. These laser strengths are similar to those used in past studies^14, 18^ and did not result in epileptiform activity in cortex.

### Histology

At the end of the experiments, all mice were transcardially perfused and tissue was processed to confirm ArchT expression was specific to PV cell populations. Briefly, mice were perfused with 30 mL 0.01M phosphate buffered saline (Fisher Scientific, BP2944-100, Pittsburgh, PA), followed by 30 mL 4% paraformaldehyde (Sigma Aldrich, 158127, St. Louis, MO). Brains were carefully removed and post-fixed 4-12 hours in 4% paraformaldehyde before being transferred to a 30% sucrose solution for at least 24 hours before sectioning. Brains were sectioned coronally at a thickness of 50µm with a freezing microtome (CM 2000R; Leica) or cryostat (CM 3050S; Leica). Tissue sections were collected throughout the auditory cortex. A subset of sections (2 sections per animal) were stained with antibodies against PV (guinea pig anti-PV antibody, SWANT GP72 1:1000) followed by Alexa Fluor 568 goat anti-guinea pig secondary antibody (No: A-11075, Thermo Fisher Scientific, 1:500). Antibodies and dilution concentrations were previously reported^67–69^. Briefly, sections were rinsed with 0.01M PBS followed by a solution of 100mM glycine (No: G7126, Sigma-Aldrich) and 0.5% Triton-X in 0.01M PBS. This was followed by a 2-hour blocking buffer incubation with 5% normal goat serum and 0.5% Triton-X in 0.01M PBS. Sections were then incubated for 24 hours with primary antibody, rinsed with 100mM glycine and 0.5% Triton-X in 0.01M PBS, and incubated with secondary antibody for 2 hours. Slices were lastly incubated for 10 min with Hoechst 33342 (No: 62249, Thermo Fisher Scientific, 1:10,000 in 0.01M PBS), rinsed with 100mM glycine and 0.5% Triton-X in 0.01M PBS before being rinsed in 100mM glycine in 0.01M PBS before mounting. Slices were mounted on slides (Fisherbrand Superfrost Plus, No: 12-5550-15, Fisher Scientific) using anti-fade mounting medium (ProLong Diamond, No: P36965, Thermo Fisher Scientific).

### Imaging and quantification

Images were taken on a VS120 widefield Olympus microscope or an OlympusFV3000 scanning confocal microscope using a 20× objective. All images were comprised of Z-stacks consisting of 5-6 slices taken at 10μm intervals throughout the 50µm slices. Stacks were taken from coronal sections as near as possible to the electrode location in auditory cortex. Areas were chosen to include similarly dense Arch-GFP cell counts across animals. To confirm targeting specificity, each PV+ cell was categorized as co-expressing or not expressing Arch-GFP across a 300×300µm grid. We also quantified the number of Arch+ cells from each stack that were not PV+ based on Hoechst labeling to estimate off target expression. We analyzed 2-4 non-overlapping stacks from 2 slices per animal from the animals that made up optogenetic Arch+ dataset (*n* = 9 PV-Arch). Cell counts were pooled across slices stained for the same marker for each animal and averaged to produce a single data point for quantification.

### Spike extraction and clustering

Kilosort2 (https://github.com/MouseLand/Kilosort) was used to detect multi-units within the recordings^26^. Before spike detection and sorting, the broadband signal was band-passed between 300 and 5000 Hz using a 3^rd^-order Butterworth filter. Kilosort results were then loaded onto Phy2 (https://github.com/cortex-lab/phy) to manually determine if spike clusters exhibited neural activity or noise^27^. Clusters with either artifact-like waveforms from laser or similar responses across all channels were deemed as noise, and spikes with artifact-like waveforms were removed from clusters that clearly exhibited neural activity, whenever possible. Clusters were merged if the cross-correlograms were similar to the component clusters’ auto-correlograms and showed overlap in principal component feature space at the same channel. The spikes toolbox (https://github.com/cortex-lab/spikes) was then used to import the cluster information from Phy to Matlab and extract spike waveforms from the high-passed signal^26^. Clusters were assigned to recording channels based on which site yielded the largest average spike amplitude. To remove any remaining artifacts from laser onset and offset, all spikes with waveforms above an absolute threshold of 1500 µV or a positive value above 750 µV were discarded, and clusters that still showed a high amount of remaining artifact after removal were excluded from further analysis. To determine which of the remaining clusters were single-units (SU), we utilized the sortingQuality toolbox (https://github.com/cortex-lab/sortingQuality) to calculate isolation distances and L-ratios^70^. SUs must 1) have less than 5% of inter-spike intervals below 2ms (Figure 1C), 2) an isolation distance above 15, and 3) an L-ratio below 0.25. For clusters where isolation distance and L-ratio were not defined, the first threshold was used. These thresholds are consistent with values used in past studies on single-unit activity^71–73^, and clusters that did not meet any of these criteria were deemed multi-units (MUs). Finally, SUs were classified as narrow-spiking if the trough-peak interval of their mean waveform was below 0.5ms, a threshold that is consistent with past findings on excitatory and inhibitory units in mouse auditory cortex^12^.

### Neural discriminability performance using SPIKE-distance

Neural discrimination performance refers to the ability to determine stimulus identity based on neural responses, thus measuring a neuron’s ability to encode stimulus features. Here, performance was calculated using a template-matching approach similar to our previous studies^25^. Briefly, spike trains were classified to one of the two target stimuli based on whose template, one from each stimulus, yielded a smaller spike distance. For each target-masker configuration, 100 iterations of template matching were done. In each iteration, one of the 10 spike trains for each target was chosen as a template, and all remaining trials were matched to each template to determine target identity. All possible pairs of templates were used across the 100 iterations to calculate an average value of neural discriminability. SPIKE-distance^21^ calculates the dissimilarity between two spike trains based on differences in spike timing and instantaneous firing rate without additional parameters. For one spike train in a pair, the instantaneous spike timing difference at time *t* is:

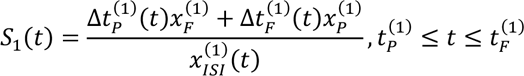

where *Δt_P_* represents the distance between the preceding spike from train 1 (*t_P_*(1)) and the nearest spike from train 2, *Δt_F_* represents the distance between the following spike from train 1 (*t_F_*(1)) and the nearest spike from train 2, *x_F_* is the absolute difference between *t* and *t_F_*(1), and *x_P_* is the absolute difference between *t* and *t_P_*(1). To calculate *S_2_(t)*, the spike timing difference from the view of the other train, all spike times and ISIs are replaced with the relevant values in train 2. The pairwise instantaneous difference between the two trains is calculated as:

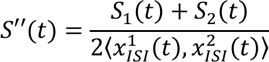

Finally, *S_1_(t)* and *S_2_(t)* are locally weighted by their instantaneous spike rates to account for differences in firing rate:

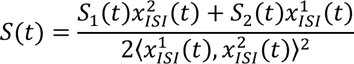

For a train of length *T*, the distance is the integral of the dissimilarity profile across the entire response interval, with a minimum value of 0 for identical spike trains:

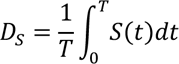

cSPIKE, a toolbox used to calculate SPIKE-distance, was used to calculate all spike train distances between all possible spike train pairs for all spatial grid configurations^21^.

To determine how firing rate modulation, spike timing, and average firing rate contribute to discriminability, we used different distance measures as inputs to the classifier. For all hotspots, performances using inter-spike interval (ISI)-distance, rate-independent (RI)-SPIKE-distance, and spike count distance, the absolute difference in spike count between trains, were also calculated and compared to SPIKE-distance-based values.

### ISI-distance

To determine how optogenetic suppression affects rapid temporal modulations in firing rate, ISI-distances were calculated. The ISI-distance calculates the dissimilarity between two spike trains based on differences in instantaneous rate synchrony. For a given time point:

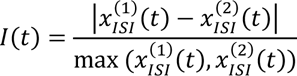

This profile is then integrated along the spike train length to give a distance value, with values of 0 obtained for either identical spike trains or pairs with the same constant firing rate and a global phase shift difference.

### RI-SPIKE-distance

To determine how optogenetic suppression affects spike timing, RI-SPIKE-distances between spike trains were calculated. The RI-SPIKE-distance is rate-independent, as it does not take differences in local firing rate between the two spike trains into account. From SPIKE-distance calculations, the final step of weighing *S_1_(t)* and *S_2_(t)* by their instantaneous spike rates is skipped, yielding:

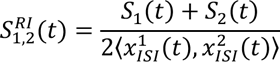

Like the other measures, the dissimilarity profile is integrated to give a distance value, with a value of 0 obtained for two identical spike trains.

### Rate-normalized RMS difference and trial similarity

In addition to average firing rate, we also calculated two other measures to determine their impact on classification performance: the similarity of responses within target, and the dissimilarity of responses across targets. To quantify intertrial reliability of responses to target stimuli, we adopted the measure of trial similarity from previous studies^35^. Specifically, we randomly divided the 10 trials in each configuration into two equal groups, binned spike times with a time resolution of 25ms, and calculated the Pearson’s correlation coefficient between the two resulting PSTHs. This process was repeated 100 times to obtain a mean correlation coefficient, or trial similarity.

We also calculated the rate-normalized RMS difference between target responses to quantify the dissimilarity in the temporal pattern of responses between the two targets. We first binned each target response using the same time-resolution as trial similarity (25ms) and normalized each PSTH such that the sum of all bins over time was 1. The RMS difference between the two rate-normalized PSTHs was then calculated. This measure quantifies the dissimilarity in the temporal pattern of responses across the targets, accounting for differences in mean evoked firing rate between targets.

All three response measures (average firing rate, trial similarity, and rate-normalized RMS difference between targets) were correlated with SPIKE-distance-based performance using Pearson’s correlation coefficients, with separate calculations done for control and laser trials.

### Decoding time analysis using van Rossum distances

To estimate the decoding time of the spike trains at each hotspot, we used van Rossum distances^36^. Briefly, the van Rossum distance between two spike trains involves convolving each response with a decaying exponential kernel with time constant *τ*. The distance between two smoothed spike trains *f_1_(t)* and *f_2_(t)* is calculated as:

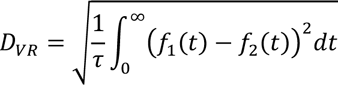

For each spatial grid configuration, a distance matrix containing the van Rossum distances between all possible spike train pairs was set as the input for the template-matching approach. Performance was calculated across a range of *τ* values, increasing in powers of 2 from 1ms to 256ms. Finally, to determine the optimal *τ* value at which performance was maximized for each configuration, we implemented a fine-grain parameter where *τ* was varied in steps of 1ms, with the optimization separately done for control and laser trials.

### Statistics and reproducibility

All single-units and spatial grid data were extracted from *N* = 9 mice. Spatial grid hotspots of high neural discriminability were determined using three criteria: 1) mean performance must be above 70% during control trials; 2) mean control performance distribution must be significantly different from chance (*p* < 0.05), calculated using a null distribution obtained by classifying spike trains within each target, which should result in chance performance; and 3) the effect size given by Cohen’s *d* between the two distributions (control vs. null) must be greater than 1:

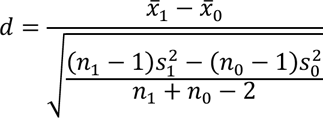

where values with subscript 0 represent the mean, standard deviation, and number of template-matching iterations for the null performance distribution. Additionally, configurations where at least 3 trials for one target showed zero spiking were excluded from analysis, to avoid inaccurate estimates of performance. This resulted in *n* = 49 clean configurations and *n* = 20 masked configurations, both of which were used to analyze the effects of suppression on discriminability and spiking activity. In the manuscript, we focus on SUs with hotspots in the control condition. We found a small number of emergent hotspots (12 from 10 single-units across both clean and masked trials) where performance and effect size were both below threshold in the control condition but above threshold in the laser condition, with a median performance 72.9% and an inter-quartile range of 2.35%. To analyze the effects of suppression on performance metrics, we used built-in Matlab functions to run paired *t*-tests between control and optogenetic values to determine statistical significance (*p*≤ 0.05), with tests done separately for clean and masked trials.

We also analyzed 47 low performance hotspots—configurations with performances between chance and our threshold of 70%—in three separate groups: hotspots with effect sizes 1) between 0.2 and 0.5, 2) between 0.5 and 0.8, and 3) greater than 0.8. To determine whether low-performance hotspots showed similar changes in performance to our main set of hotspots, we ran paired t-tests and calculated the effect size of optogenetic suppression on discriminability. For the first group, we found 68 clean configurations and 352 masked configurations. Clean performance did not significantly decrease (*p* = 0.8116, *d* = −0.03) while masked performance did (*p* < 1e-04, *d* = 0.24). For the second group, both clean (*n* = 56, *p* = 0.0150, *d* = 0.34) and masked (*n* = 166, *p* < 1e-04, *d* = 0.67) performance decreased with suppression. For the last group, both clean (*n* = 35, *p* = 0.0043, *d* = 0.52) and masked (*n* = 98, *p* < 1e-04, *d* = 0.84) performance decreased with suppression.

## Supporting information

Supplementary analysis and results

## Acknowledgments

This research was supported by the National Science Foundation (#IIS-1835270), the National Institute of Health (#1R34NS111742-01 and #1T32DC013017-01A1), and the Boston University Micro and Nano Imaging Facility (NIH S10OD024993) We would like to thank Alberto Cruz-Martín, Michael Economo, and Conor Houghton for comments and suggestions on the manuscript. We also thank Monty Escabi and Oded Ghitza for discussions on coding time-scales in auditory cortex and speech.

## Competing Interest Statement

The authors declare no conflicts of interest.

## Data Availability Statement

The data used for the analysis in this study is available upon request. The source data for the figures and code used in this study are available at github.com/NSNC-Lab/SpatialDiscriminabilityAnalysis.

## Notes

### Competing Interest Statement

The authors have declared no competing interest.

### Summary of Updates

L-ratio and isolation distance added as additional metrics to single-unit classification; new supplemental analysis of opto-genetic suppression of PV+ cells; new supplemental results from non-Arch-expressing mice to rule out non-specific effects of illumination; histological analysis of Arch expression in experimental group; statistics and reproducibility section added to Methods; Figures and supplemental data revised to reflect refined single-unit results; authors responsible for histology and control experiments added

## References

1. deCharms RC, Zador A. Neural representation and the cortical code. Annu Rev Neurosci 23, 613–647 (2000).

2. Zuo Y, Safaai H, Notaro G, Mazzoni A, Panzeri S, Diamond ME. Complementary contributions of spike timing and spike rate to perceptual decisions in rat S1 and S2 cortex. Curr Biol 25, 357–363 (2015).

3. Yao JD, Sanes DH. Temporal Encoding is Required for Categorization, But Not Discrimination. Cereb Cortex 31, 2886–2897 (2021).

4. Lu T, Liang L, Wang X. Temporal and rate representations of time-varying signals in the auditory cortex of awake primates. Nat Neurosci 4, 1131–1138 (2001).

5. Tremblay R, Lee S, Rudy B. GABAergic Interneurons in the Neocortex: From Cellular Properties to Circuits. Neuron 91, 260–292 (2016).

6. Pfeffer CK, Xue M, He M, Huang ZJ, Scanziani M. Inhibition of inhibition in visual cortex: the logic of connections between molecularly distinct interneurons. Nat Neurosci 16, 1068–1076 (2013).

7. Canales A, Scheuer KS, Zhao X, Jackson MB. Unitary synaptic responses of parvalbumin interneurons evoked by excitatory neurons in the mouse barrel cortex. Cereb Cortex, (2022).

8. Antonoudiou P, Tan YL, Kontou G, Upton AL, Mann EO. Parvalbumin and Somatostatin Interneurons Contribute to the Generation of Hippocampal Gamma Oscillations. J Neurosci 40, 7668–7687 (2020).

9. Jang HJ, Chung H, Rowland JM, Richards BA, Kohl MM, Kwag J. Distinct roles of parvalbumin and somatostatin interneurons in gating the synchronization of spike times in the neocortex. Sci Adv 6, eaay5333 (2020).

10. Wehr M, Zador AM. Balanced inhibition underlies tuning and sharpens spike timing in auditory cortex. Nature 426, 442–446 (2003).

11. Li LY, et al. A feedforward inhibitory circuit mediates lateral refinement of sensory representation in upper layer 2/3 of mouse primary auditory cortex. J Neurosci 34, 13670–13683 (2014).

12. Li LY, Xiong XR, Ibrahim LA, Yuan W, Tao HW, Zhang LI. Differential Receptive Field Properties of Parvalbumin and Somatostatin Inhibitory Neurons in Mouse Auditory Cortex. Cereb Cortex 25, 1782–1791 (2015).

13. Moore AK, Wehr M. Parvalbumin-expressing inhibitory interneurons in auditory cortex are well-tuned for frequency. J Neurosci 33, 13713–13723 (2013).

14. Aizenberg M, Mwilambwe-Tshilobo L, Briguglio JJ, Natan RG, Geffen MN. Bidirectional Regulation of Innate and Learned Behaviors That Rely on Frequency Discrimination by Cortical Inhibitory Neurons. PLoS Biol 13, e1002308 (2015).

15. Natan RG, et al. Complementary control of sensory adaptation by two types of cortical interneurons. Elife 4, (2015).

16. Blackwell JM, Geffen MN. Progress and challenges for understanding the function of cortical microcircuits in auditory processing. Nat Commun 8, 2165 (2017).

17. Seybold BA, Phillips EAK, Schreiner CE, Hasenstaub AR. Inhibitory Actions Unified by Network Integration. Neuron 87, 1181–1192 (2015).

18. Phillips EA, Hasenstaub AR. Asymmetric effects of activating and inactivating cortical interneurons. Elife 5, (2016).

19. Moore AK, Weible AP, Balmer TS, Trussell LO, Wehr M. Rapid Rebalancing of Excitation and Inhibition by Cortical Circuitry. Neuron 97, 1341–1355 e1346 (2018).

20. Penikis KB, Sanes DH. A Redundant Cortical Code for Speech Envelope. J Neurosci 43, 93–112 (2023).

21. Satuvuori E, Kreuz T. Which spike train distance is most suitable for distinguishing rate and temporal coding? J Neurosci Methods 299, 22–33 (2018).

22. Kreuz T, Chicharro D, Houghton C, Andrzejak RG, Mormann F. Monitoring spike train synchrony. J Neurophysiol 109, 1457–1472 (2013).

23. Narayan R, et al. Cortical interference effects in the cocktail party problem. Nat Neurosci 10, 1601–1607 (2007).

24. Mesgarani N, Chang EF. Selective cortical representation of attended speaker in multi-talker speech perception. Nature 485, 233–236 (2012).

25. Maddox RK, Billimoria CP, Perrone BP, Shinn-Cunningham BG, Sen K. Competing sound sources reveal spatial effects in cortical processing. PLoS Biol 10, e1001319 (2012).

26. Pachitariu M, Steinmetz N, Kadir S, Carandini M, Harris KD. Kilosort: realtime spike-sorting for extracellular electrophysiology with hundreds of channels. bioRxiv, (2016).

27. Rossant C, et al. Spike sorting for large, dense electrode arrays. Nat Neurosci 19, 634–641 (2016).

28. Keller CH, Kaylegian K, Wehr M. Gap encoding by parvalbumin-expressing interneurons in auditory cortex. J Neurophysiol 120, 105–114 (2018).

29. Rothauser EH, Chapman, W. D., Guttman, N., Nordby, K. S., Silbiger, H. R., Urbanek, G. E., & Weinstock, M. I.E.E.E. recommended practice for speech quality measurements. IEEE Trans Audio Electroacoust 17, 225–246 (1969).

30. Britten KH, Shadlen MN, Newsome WT, Movshon JA. The analysis of visual motion: a comparison of neuronal and psychophysical performance. J Neurosci 12, 4745–4765 (1992).

31. Parker AJ, Newsome WT. Sense and the single neuron: probing the physiology of perception. Annu Rev Neurosci 21, 227–277 (1998).

32. Wang L, Narayan R, Grana G, Shamir M, Sen K. Cortical discrimination of complex natural stimuli: can single neurons match behavior? J Neurosci 27, 582–589 (2007).

33. Billimoria CP, Kraus BJ, Narayan R, Maddox RK, Sen K. Invariance and sensitivity to intensity in neural discrimination of natural sounds. J Neurosci 28, 6304–6308 (2008).

34. Downer JD, Bigelow J, Runfeldt MJ, Malone BJ. Temporally precise population coding of dynamic sounds by auditory cortex. J Neurophysiol 126, 148–169 (2021).

35. Hoglen NEG, Larimer P, Phillips EAK, Malone BJ, Hasenstaub AR. Amplitude modulation coding in awake mice and squirrel monkeys. J Neurophysiol 119, 1753–1766 (2018).

36. van Rossum MC. A novel spike distance. Neural Comput 13, 751–763 (2001).

37. Scala F, et al. Phenotypic variation of transcriptomic cell types in mouse motor cortex. Nature, (2020).

38. Cardin JA, et al. Driving fast-spiking cells induces gamma rhythm and controls sensory responses. Nature 459, 663–667 (2009).

39. Sohal VS, Zhang F, Yizhar O, Deisseroth K. Parvalbumin neurons and gamma rhythms enhance cortical circuit performance. Nature 459, 698–702 (2009).

40. Bruno RM, Simons DJ. Feedforward mechanisms of excitatory and inhibitory cortical receptive fields. J Neurosci 22, 10966–10975 (2002).

41. Xiang Z, Huguenard JR, Prince DA. Cholinergic switching within neocortical inhibitory networks. Science 281, 985–988 (1998).

42. Wang JA, McFadden SL, Caspary D, Salvi R. Gamma-aminobutyric acid circuits shape response properties of auditory cortex neurons. Brain Research 944, 219–231 (2002).

43. Kurt S, Moeller CK, Jeschke M, Schulze H. Differential effects of iontophoretic application of the GABAA-antagonists bicuculline and gabazine on tone-evoked local field potentials in primary auditory cortex: interaction with ketamine anesthesia. Brain Res 1220, 58–69 (2008).

44. Chen QC, Jen PH. Bicuculline application affects discharge patterns, rate-intensity functions, and frequency tuning characteristics of bat auditory cortical neurons. Hear Res 150, 161–174 (2000).

45. Kato HK, Asinof SK, Isaacson JS. Network-Level Control of Frequency Tuning in Auditory Cortex. Neuron 95, 412–423 e414 (2017).

46. Briguglio JJ, Aizenberg M, Balasubramanian V, Geffen MN. Cortical Neural Activity Predicts Sensory Acuity Under Optogenetic Manipulation. J Neurosci 38, 2094–2105 (2018).

47. Liang F, et al. Sparse Representation in Awake Auditory Cortex: Cell-type Dependence, Synaptic Mechanisms, Developmental Emergence, and Modulation. Cereb Cortex 29, 3796–3812 (2019).

48. Theunissen F, Miller JP. Temporal encoding in nervous systems: a rigorous definition. J Comput Neurosci 2, 149–162 (1995).

49. Ding N, Patel AD, Chen L, Butler H, Luo C, Poeppel D. Temporal modulations in speech and music. Neurosci Biobehav Rev 81, 181–187 (2017).

50. Castellucci GA, Calbick D, McCormick D. The temporal organization of mouse ultrasonic vocalizations. PLoS One 13, e0199929 (2018).

51. Chen C, Read HL, Escabi MA. Precise feature based time scales and frequency decorrelation lead to a sparse auditory code. J Neurosci 32, 8454–8468 (2012).

52. Ghitza O. Linking speech perception and neurophysiology: speech decoding guided by cascaded oscillators locked to the input rhythm. Front Psychol 2, 130 (2011).

53. Teng X, Poeppel D. Theta and Gamma Bands Encode Acoustic Dynamics over Wide-Ranging Timescales. Cereb Cortex 30, 2600–2614 (2020).

54. Shadlen MN, Newsome WT. Noise, neural codes and cortical organization. Curr Opin Neurobiol 4, 569–579 (1994).

55. Narayan R, Ergun A, Sen K. Delayed inhibition in cortical receptive fields and the discrimination of complex stimuli. J Neurophysiol 94, 2970–2975 (2005).

56. Dong J, Colburn HS, Sen K. Cortical Transformation of Spatial Processing for Solving the Cocktail Party Problem: A Computational Model(1,2,3). eNeuro 3, (2016).

57. Kapfer C, Glickfeld LL, Atallah BV, Scanziani M. Supralinear increase of recurrent inhibition during sparse activity in the somatosensory cortex. Nat Neurosci 10, 743–753 (2007).

58. Silberberg G, Markram H. Disynaptic inhibition between neocortical pyramidal cells mediated by Martinotti cells. Neuron 53, 735–746 (2007).

59. Adesnik H, Bruns W, Taniguchi H, Huang ZJ, Scanziani M. A neural circuit for spatial summation in visual cortex. Nature 490, 226–231 (2012).

60. Lakunina AA, Nardoci MB, Ahmadian Y, Jaramillo S. Somatostatin-Expressing Interneurons in the Auditory Cortex Mediate Sustained Suppression by Spectral Surround. J Neurosci 40, 3564–3575 (2020).

61. Norman-Haignere SV, et al. Multiscale temporal integration organizes hierarchical computation in human auditory cortex. Nat Hum Behav 6, 455–469 (2022).

62. Chou KF, Sen K. AIM: A network model of attention in auditory cortex. PLoS Comput Biol 17, e1009356 (2021).

63. Gritton HJ, et al. Oscillatory activity in alpha/beta frequencies coordinates auditory and prefrontal cortices during extinction learning. bioRxiv, (2020).

64. James NM, Gritton HJ, Kopell N, Sen K, Han X. Muscarinic receptors regulate auditory and prefrontal cortical communication during auditory processing. Neuropharmacology 144, 155–171 (2019).

65. Hawley ML, Litovsky RY, Culling JF. The benefit of binaural hearing in a cocktail party: effect of location and type of interferer. J Acoust Soc Am 115, 833–843 (2004).

66. Zhu Y, Qiao W, Liu K, Zhong H, Yao H. Control of response reliability by parvalbumin-expressing interneurons in visual cortex. Nat Commun 6, 6802 (2015).

67. Keaveney MK, Tseng HA, Ta TL, Gritton HJ, Man HY, Han X. A MicroRNA-Based Gene-Targeting Tool for Virally Labeling Interneurons in the Rodent Cortex. Cell Rep 24, 294–303 (2018).

68. Gritton HJ, et al. Unique contributions of parvalbumin and cholinergic interneurons in organizing striatal networks during movement. Nat Neurosci 22, 586–597 (2019).

69. Tseng H-a, Mount RA, Lowet E, Gritton HJ, Cheung C, Han X. Membrane Voltage Dynamics of Parvalbumin Interneurons Orchestrate Hippocampal Theta Rhythmicity. bioRxiv, 2022.2011.2014.516448 (2022).

70. Schmitzer-Torbert N, Jackson J, Henze D, Harris K, Redish AD. Quantitative measures of cluster quality for use in extracellular recordings. Neuroscience 131, 1–11 (2005).

71. Kvitsiani D, Ranade S, Hangya B, Taniguchi H, Huang JZ, Kepecs A. Distinct behavioural and network correlates of two interneuron types in prefrontal cortex. Nature 498, 363–366 (2013).

72. Monaghan JJM, Garcia-Lazaro JA, McAlpine D, Schaette R. Hidden Hearing Loss Impacts the Neural Representation of Speech in Background Noise. Curr Biol 30, 4710–4721 e4714 (2020).

73. Jung F, Yanovsky Y, Brankack J, Tort ABL, Draguhn A. Respiratory entrainment of units in the mouse parietal cortex depends on vigilance state. Pflugers Arch, (2022).

74. Wesson DW, Donahou TN, Johnson MO, Wachowiak M. Sniffing behavior of mice during performance in odor-guided tasks. Chem Senses 33, 581–596 (2008).

