## Supplementary analysis and results for "Parvalbumin neurons, temporal coding, and cortical noise in complex scene analysis"

**Classification:** Biological Sciences/Neuroscience

**Keywords:** Parvalbumin, cortical code, temporal code, cortical noise, cocktail party problem, complex scene analysis

**Author contributions:** J.C.N. and H.J.G. performed all experiments. J.C.N. analyzed the data. N.M.J. provided technical support and code for analysis. R.A.M. performed histological analysis. Z.Q. and H.J.G. performed additional experiments for control analysis. X.H. and K.S obtained funding and supervised the study. J.C.N., H.J.G., X.H., and K.S. wrote the manuscript and contributed to the interpretation of the results.

### Supplemental Methods

**Current source density estimation and layer analysis**. Current source density (CSD) analysis estimates the second spatial derivative of LFP signals to determine the relative current across the cortical laminar depth. CSDs were calculated using LFPs, as described previously[1]. LFPs from control masked trials were used, as the rise time was more similar between target identities than clean stimuli. LFPs were low-passed filtered at 150 Hz before being down-sampled by a factor of 8 to 381 Hz. For each channel, LFPs were averaged across all control masked trials prior to CSD estimation, and channels that did not show an evoked response were interpolated using neighboring sites on the same shank. After this, LFPs were spatial smoothed across the eight channels in each shank:

$$\phi(z) = \frac{\phi(z + \Delta z) + 2\phi(z) + \phi(z - \Delta z)}{4}$$

where z is the depth perpendicular to the cortical surface, $\Delta$z is the electrode spacing, and $\Phi$ is the potential. CSD was then estimated as:

$$CSD(z) = -\frac{\phi(z + \Delta z) - 2\phi(z) + \phi(z - \Delta z)}{\Delta z^2}$$

To determine the granular layer in each shank, CSD sink onset times were calculated as the time when the CSD goes below -3 times the standard deviation of pre-stimulus activity. If more than one channel was found to have the earliest sink onset, the channel whose neighbors had the earliest onsets was deemed the granular layer, or L4. The width of each layer was estimated based on previous anatomical studies[2]. L1 consisted of channels at least 500μm above the input layer, L2/3 consisted of channels 200μm to 400μm above the channel with the earliest sink onset; L4 consisted of the input channel and the channel 100μm above it; L5 consisted of channels 100 to 300μm below the input layer, and L6 consisted of all channels at 400μm below the input layer.

**Statistical analysis.** For performance comparisons between layers and between narrow-spiking and regular-spiking units, we separately ran repeated-measures ANOVA and effect size calculations for clean and masked trials, with condition as the within-subjects factor and cell type (narrow-spiking or regular-spiking) or layer as the between-subjects factor. Effect sizes were calculated using the Measures of Effect Size toolbox[3, 4], and post-hoc Tukey-Kramer tests were carried out if the ANOVA returned significance. To estimate the effect of optogenetic suppression on spiking across layers, repeated-measures ANOVA was carried out for both spontaneous and onset firing rate. Spontaneous firing rate was defined as the average firing rate during the 50ms between laser onset and sound onset, while onset firing rate was defined as the average rate during the first 0.5s of sound stimulus playback. For both measures, ANOVA was done for regular-spiking single units, with layer as the between-subjects factor and condition as the within-subjects factor. Post-hoc Tukey-Kramer tests were carried out if the ANOVA returned a significant factor or interaction. Repeated-measures ANOVA was not done for narrow-spiking single units due to the small sample size per layer[5].

### Supplemental References

1. James NM, Gritton HJ, Kopell N, Sen K, Han X. Muscarinic receptors regulate auditory and prefrontal cortical communication during auditory processing. *Neuropharmacology* **144**, 155-171 (2019).
2. Morrill RJ, Hasenstaub AR. Visual Information Present in Infragranular Layers of Mouse Auditory Cortex. *J Neurosci* **38**, 2854-2862 (2018).

3. Hentschke H, Stuttgen MC. Computation of measures of effect size for neuroscience data sets. *Eur J Neurosci* **34**, 1887-1894 (2011).
4. Hentschke H. hhentschke/measures-of-effect-size-toolbox.) (2022).
5. Serdar CC, Cihan M, Yucel D, Serdar MA. Sample size, power and effect size revisited: simplified and practical approaches in pre-clinical, clinical and laboratory studies. *Biochem Med (Zagreb)* **31**, 010502 (2021).

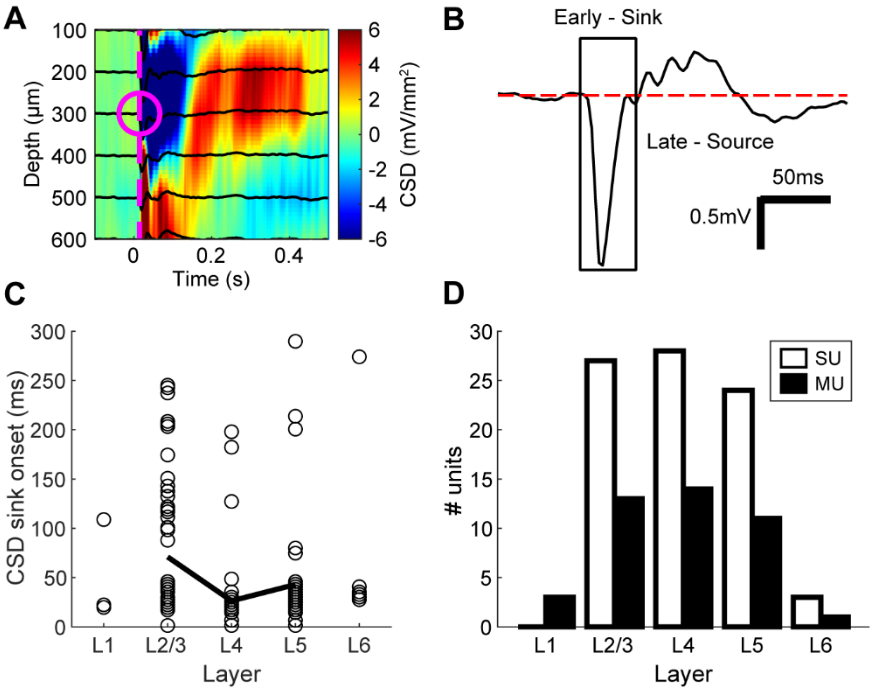


**Supplementary Figure 1. Recording locations and density using CSD for layer separation.** **A:** Current source density pseudo-colormap with event-related potential (ERP) traces overlaid in black. Magenta dashed line indicates time of earliest CSD sink onset, while the magenta circle indicates the channel with earliest onset. The mean CSD sink onset at all identified granular channels was 17.2 ms. **B:** Example ERP trace with CSD sink boxed in. Red dashed line indicates mean pre-stimulus activity, which was used to determine the threshold below which the CSD sink was detected. **C:** Layer vs. CSD sink onsets for all channels, with mean sink onset outlined in black from L2/3 to L5, with the mean CSD sink onset at L4 at 26.4ms. Mean sink onset for L1 and L6 are not shown due to the lack of units in both layers. **D:** Bar plot showing number of detected single units (SU, white, n = 82 units) and multi-units (MU, black, n = 42 units) at each layer.

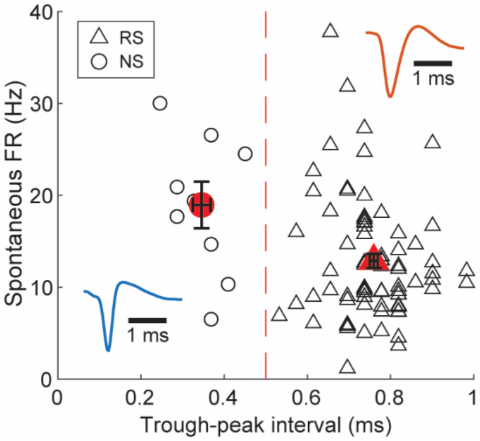


**Supplementary Figure 2. Single units with narrow-spiking waveforms vs. regular-spiking waveforms.** Scatter plot showing trough-peak interval of each single unit's spike waveform versus spontaneous firing rate during control trials. Dashed red line at 0.5ms represents the threshold between narrow-spiking units (NS, circle markers, n = 9 units) and regular-spiking units (RS, triangle markers, n = 73 units). Insets show example waveforms from a narrow-spiking single unit (bottom-left, blue) and regular-spiking single unit (top-right, orange) with scale bars measuring 1ms. Filled red markers represent the mean trough-peak interval and spontaneous firing rate for each unit type, with error bars representing ± SEM. Using two-sample t-tests with an assumption of unequal variance, both trough-peak interval ($p < 1e\text{-}04$, $d = 4.42$) and spontaneous firing rate ($p = 0.0460$, $d = -0.89$) were found to be significantly different between NS and RS units.

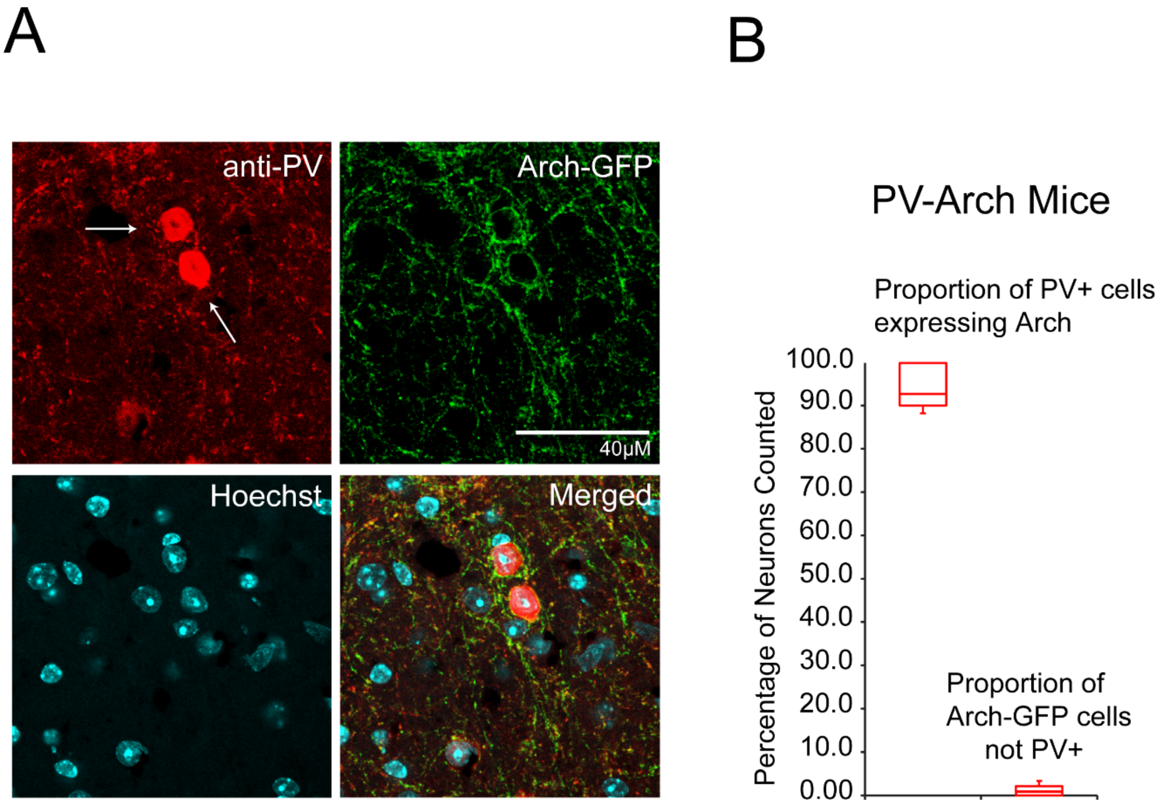


**Supplementary Figure 3. Histology of PV-Arch mice: A:** Post-study representative confocal photomicrograph example from PV-Arch mouse used in this study. Example includes merged images along with separate channel images for Arch-GFP (green), anti-PV immunofluorescence (red), and Hoechst labeling (cyan). White arrows indicate cells with co-localized expression of Arch and anti-PV. **B**: Box plot indicating quantification of viral specificity from auditory regions in PV-Arch mice recorded in this study (*n* = 9; see 'Methods' in main text). 93.5% ± 1.0% (mean ± s.e.m.) of PV immunoreactive cells co-expressed Arch-GFP while 0.95% ± 0.23% (mean ± s.e.m.) of Arch-GFP cells were not immunoreactive for PV. For box plot figures, middle lines indicate the median, lower and upper lines of the box indicate quartiles below and above the median, and upper and lower whiskers indicate maximum or minimum values respectively.

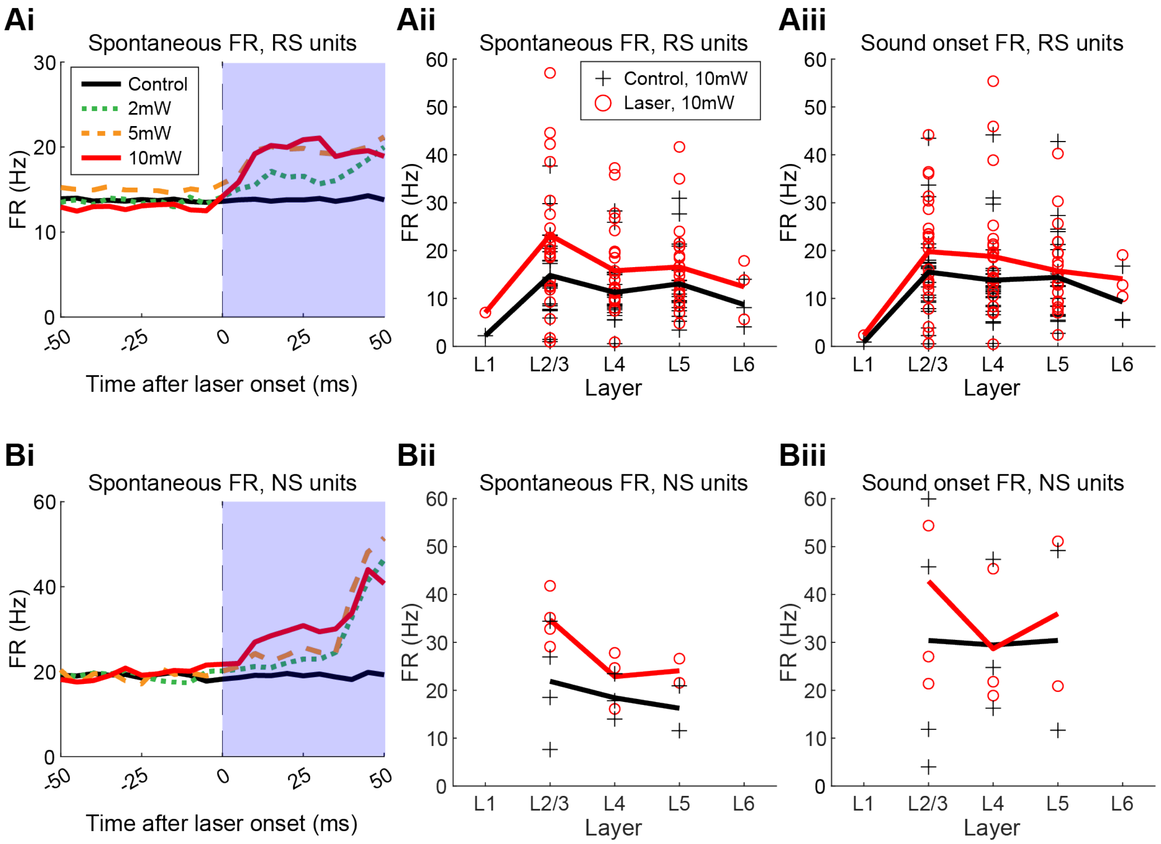


**Supplementary Figure 4. Effects of optogenetic suppression on firing rate across layers for regular-spiking (RS) and narrow-spiking (NS) single units. Ai:** Traces of average spontaneous firing rate from 50ms before to 50ms after laser onset (thin dashed line) for control trials (solid black line) and all laser powers (2mW, green dotted line; 5mW orange dashed line; 10mW, red solid line) for all RS units. Blue shaded region represents the time period during which laser was on, but no sound was present. During the 100ms shown, no auditory stimulus was presented. The control trace was estimated by pooling all non-optogenetic trials from each of the three blocks. Spontaneous firing rate increases with optogenetic suppression of PV neurons, in a graded manner with laser power, consistent with previous studies. **Aii:** Spontaneous firing rate for all RS units across layer (L1: n = 1 unit, L2/3: n = 24 units; L4: n = 26 units; L5: n = 22 units; L6: n = 3 units) during control and laser trials at the 10mW recording block, estimated during the 50ms-long blue region in Ai. L1 was excluded from further analysis due to the small sample size. Repeated-measures ANOVA found laser (p(3,71) = 6e-04, $\eta^2_p$ = 0.15) as a significant factor, but not layer (p(3,71) = 0.094, $\eta^2_p$ = 0.09) or the interaction between the two (p(3,71) = 0.213, $\eta^2_p$ = 0.06). Within layers, post-hoc tests found highly significant differences between conditions in L2/3 (p = 7e-03, d = -0.68) and L4 ($p < 1e-04$, d = -0.68). **Aiii:** Sound onset firing rate (the first 0.5 s of stimulus presentation across both clean and masked trials) for RS units across layer during the 10mW recording block. Repeated-measures ANOVA yielded laser as a significant factor (p(3,71) < 1e-04, $\eta^2_p$ = 0.25) but not for layer (p(3,71) = 0.692, $\eta^2_p$ = 0.02) or interaction between the two (p(3,71) = 0.05, $\eta^2_p$ = 0.10). Within layers, post-hoc tests found highly significant differences between conditions in L2/3 ($p < 1e-04$, d = -0.74) and L4 ($p < 1e-04$, d = -0.05). **Bi:** Traces of average spontaneous firing rate from 50ms before to 50ms after laser onset (thin dashed line) for control trials (solid black line) and all laser powers (2mW, green dotted line; 5mW orange dashed line; 10mW, red solid line) for all NS units. Counterintuitively, but consistent with the study by Moore et al., spontaneous firing rate for NS units also increases upon PV suppression. **Bii:** Spontaneous firing rate for all regular-spiking single units across layer (L2/3: n = 4 units; L4: n = 3 units; L5: n = 2 units) during control and laser trials at the 10mW recording block, estimated during the 50ms-long blue region in Bi. Repeated-measures ANOVA was not calculated due to the small sample size per group. **Biii:** Sound onset firing rate for narrow-spiking single units across layer during the 10mW recording block. As with the data in Bii, repeated-measures ANOVA was not calculated due to the small sample size per group.

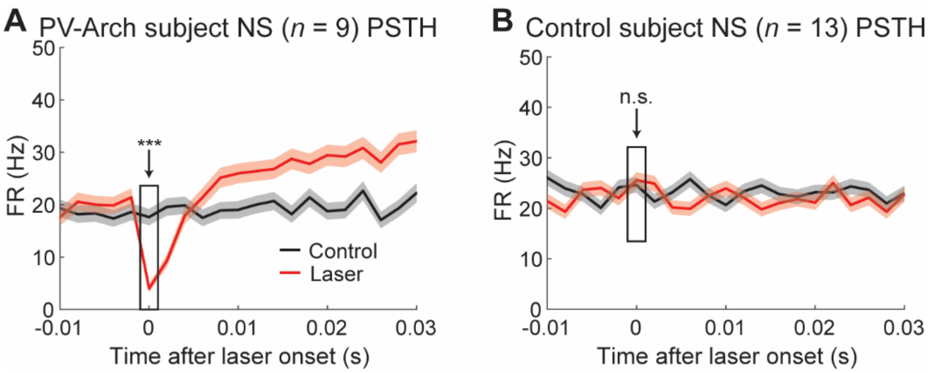


**Supplemental Figure 5. NS units in PV-Arch subjects show immediate suppression following laser onset. A:** Mean trial PSTH of all 9 PV-Arch subject NS single units around laser onset during control (black) and optogenetic (red) conditions, with a time-resolution of 2 ms and shaded regions representing ± s.e.m. at each time bin. Since no auditory stimulus is present during this period, both clean and masked trials were used to construct the PSTHs. Using a 2-sample *t*-test, we found that the mean spiking rate during the first 2 ms of optogenetic suppression (indicated by the black box and arrow) was significantly lower ($p$ < 1e-04, $d$ = 0.19) than that of the PV non-Arch control condition. Individually, 8 of the 9 NS PV-Arch units showed a significant decrease in spiking within the first 2 ms of optogenetic suppression compared to the control. **B:** Mean trial PSTH of all 13 PV-only control-subject NS single units around laser onset during control (black) and laser (red) conditions, with a time-resolution of 2 ms and shaded regions representing ± s.e.m. at each time bin. Since no auditory stimulus is present during this period, both clean and masked trials were used to construct the PSTHs. Using a 2-sample *t*-test, we found that the mean spiking rate during the first 2 ms of laser onset was not significantly different between both conditions ($p$ = 0.6942, $d$ = 0). Individually, 12 of the 13 NS PV only units did not show a significant change in spiking during the first 2 ms of laser stimulation compared to the control.

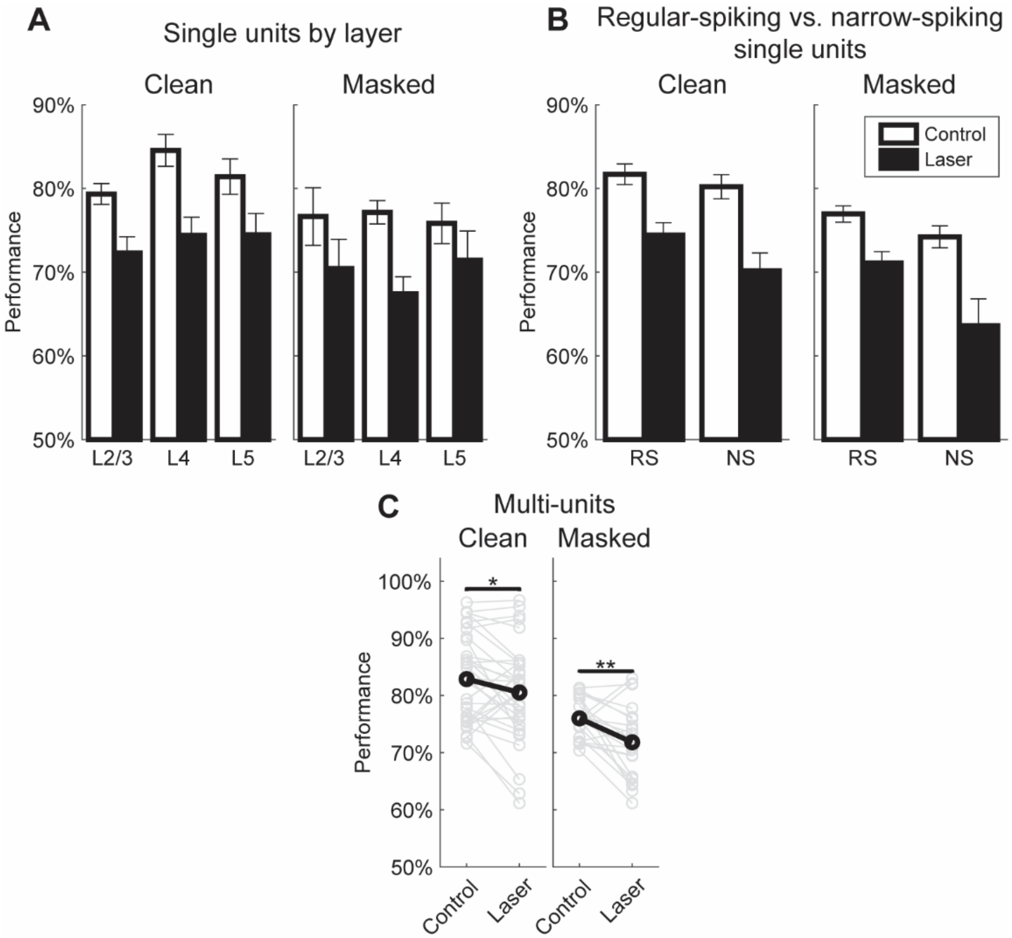


**Supplementary Figure 6. Comparison of SPIKE-distance-based performance across different unit types. A:** Comparisons between layers for all single units with hotspots (L2/3: n = 20 clean configurations, n = 3 masked configurations; L4: n = 12 clean configurations, n = 11 masked configurations; L5: n = 17 clean configurations, n = 6 masked configurations), with error bars representing SEM. Repeated-measures ANOVA for clean trials yielded significance for laser ($p(2,46) < 1e\text{-}04$, $\eta^2_p = 0.54$) but not for layer ($p(2,46) = 0.270$, $\eta^2_p = 0.06$) or interaction ($p(2,46) = 0.487$, $\eta^2_p = 0.03$). Masked ANOVA also yielded significance for laser ($p(2,17) < 1e\text{-}04$, $\eta^2_p = 0.43$) but not for layer ($p(2,17) = 0.852$, $\eta^2_p = 0.02$) or interaction ($p(2,17 = 0.360$, $\eta^2_p = 0.11$). **B:** Comparisons between regular-spiking (RS: n = 39 clean configurations, n = 17 masked configurations) and narrow-spiking (NS: n = 10 clean configurations, n = 3 masked configurations) single units. Repeated-measures ANOVA for clean trials yielded a significant effect from laser ($p(1,47) < 1e\text{-}04$, $\eta^2_p = 0.48$) but not for unit type ($p(1,47) = 0.246$, $\eta^2_p = 0.03$) or the interaction between the two ($p(1,47) = 0.268$, $\eta^2_p = 0.03$). Repeated-measures ANOVA for masked trials yielded laser as a significant factor ($p(1,18) < 1e\text{-}04$, $\eta^2_p = 0.44$) but not for unit type ($p(1,18) = 0.134$, $\eta^2_p = 0.12$) or interaction ($p(1,18) = 0.459$, $\eta^2_p = 0.03$). **C:** Comparisons between conditions for multi-units (MUs). MUs showed similar trends to SUs with a decrease in performance upon PV suppression. Paired t-tests yielded a significant decrease in performance for both clean trials (n = 33 configurations, p = 0.040, d = 0.37) and masked trials (n = 21 configurations, p = 0.0045, d = 0.70).

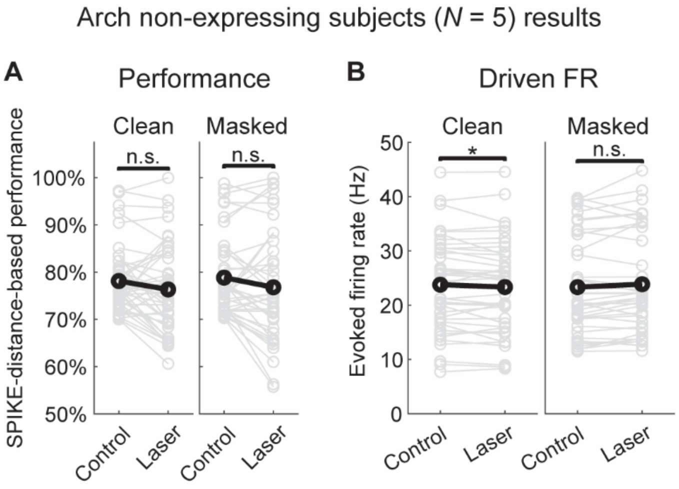


**Supplemental Figure 7. Single-units from PV-Cre -non-Arch expressing mice do not show similar changes in discriminability and spiking as PV-Arch-expressing mice. A:** Paired comparisons of SPIKE-distance-based performance from control and laser-on trials in $n$ = 5 PV non-Arch expressing mice and $n$ = 21 single units. Paired $t$-tests did not yield a significant decrease in performance for either clean ($n$ = 46 configurations; $p$ = 0.0514, $d$ = 0.30) or masked ($n$ = 40 configurations; $p$ = 0.0689, $d$ = 0.30) trials when the laser was turned on, which suggests that non-optogenetic effects from the laser did not result in the significant and strong decreases in performance (Figure 3, Clean: $p$ < 1e-04, $d$ = 1.05; Masked: $p$ < 1-e04, $d$ = 1.03) seen in our set of PV-Arch-expressing mice. **B:** Paired comparisons of mean evoked firing rate during control and laser trials. Paired $t$-tests yielded a significant change in spiking during clean trials ($p$ = 0.0482, $d$ = 0.30) but not during masked trials ($p$ = 0.2994, $d$ = -0.28).
